# A Heat Stress Responsive NAC Transcription Factor Heterodimer Plays Key Roles in Rice Grain Filling

**DOI:** 10.1101/2020.02.08.939728

**Authors:** Ye Ren, Zhouquan Huang, Hao Jiang, Zhuo Wang, Fengsheng Wu, Yufei Xiong, Jialing Yao

## Abstract

High temperature often leads to the failure of grain filling in rice (*Oryza sativa*) to cause yield loss, while the mechanism is not well elucidated yet. Here, we report that two seed-specific NAM/ATAF/CUC domain transcription factors, *ONAC127* and *ONAC129*, are responsive to heat stress and involved in the grain filling process of rice. *ONAC127* and *ONAC129* are dominantly expressed in the pericarp and can form a heterodimer during rice grain filling. CRISPR/Cas9 induced mutants and overexpression lines were then generated to investigate the functions of these two transcription factors. Interestingly, both knock-out and overexpression plants showed incomplete grain filling and shrunken grains, which became more severe under heat stress. Transcriptome analysis revealed that *ONAC127* and *ONAC129* mainly regulate stimulus response and nutrient transport. ChIP-seq analysis identified that the direct targets of ONAC127 and ONAC129 in developing rice seeds include monosaccharide transporter *OsMST6*, sugar transporter *OsSWEET4*, calmodulin-like protein *OsMSR2* and AP2/ERF factor *OsEATB*. These results suggest that *ONAC127* and *ONAC129* may regulate grain filling through affecting sugar transportation and abiotic stress responses. Overall, this study demonstrates a transcriptional regulatory network involving ONAC127 and ONAC129 and coordinating multiple pathways to modulate seed development and heat stress response at rice reproductive stage.

**Highlight:** A NAC transcription factor heterodimer plays vital roles in heat stress response and sugar transportation at rice grain filling stage.

## Introduction

Rice (*Oryza sativa*) seed development initiates from a fertilized ovary and ends with a dehydrated, hard and transparent grain, which can be divided into three stages including cell division, organogenesis and maturation (Agarwal *et al.*, 2011). A rice seed is mainly composed of endosperm with abundant starch grains, and starch biosynthesis is closely related to the transport of carbohydrate from leaves to seeds via the phloem (Patrick, 1997). The dorsal vascular bundles that pass through the pericarp are the main nutrient transport tissue in rice seeds (Oparka and Gates, 1981), while they are not connected to the endosperm (Hoshikawa, 1984). Hence, the apoplasmic pathway is the only channel for nutrients to reach the starchy endosperm (Matsuda *et al.*, 1979).

Carbohydrates may directly enter the nucellar epidermis through plasmodesmata, and then be transported to the apoplasmic space under the action of Sugar Will Eventually Be Exported Transporters (OsSWEETs) and partially hydrolyzed into monosaccharides by cell wall invertase OsCINs. The monosaccharides are then transported into the aleurone layer mainly by monosaccharide transporter OsMSTs, while the rest un-hydrolyzed sucrose is directly transported into the aleurone layer by the sucrose transporter OsSUTs (Yang *et al.*, 2018). The nutrient transport is also regulated by a range of transcription factors (TFs) such as *OsNF-YB1* and *OsNF-YC12*, which activate the expression of OsSUTs to capture the leaked sucrose in the apoplasmic space. Knockout of *NF-YB1* or *OsNF-YC12* led to defective grains with chalky endosperms, and a similar phenotype was also observed in the mutant of *OsSUT1* (Bai *et al.*, 2016, Xiong *et al.*, 2019).

The nutrient transport processes in the seed are highly susceptible to variations in environmental conditions, and extreme external stimuli (especially extreme temperature) during the grain-filling stage can lead to a nearly 50% reduction in rice yield (Hu and Xiong, 2014). Plants respond to unexpectedly high temperature through some stress-specific signaling pathways. During heat stress signal transduction, the heat shock transcription factors (HSFs) will be activated to regulate the expression of downstream heat shock proteins (HSPs) and other stress-related genes. For example, AP2/EREBP TF *DREB2A* can activate a heat shock TF *hsfA3* under heat stress to induce the expression of specific HSPs (Schramm *et al.*, 2008). Similarly, the expression of *OsbZIP60*, which plays important roles in moisture retention and heat-damage resistance, is induced under heat stress and endoplasmic reticulum stress (Oono *et al.*, 2010). Although several proteins related to plant heat stress response have been found and characterized, the mechanisms of the heat stress response are still poorly understood.

NAC (NAM, ATAF1/2, CUC2) TFs are one of the largest family of plant specific TFs with 151 members in rice (Nuruzzaman *et al.*, 2010). The typical structure of NAC TFs is a conserved NAC domain with about 150 amino acids in the N-terminus, and a variable transcriptional regulation region in the C-terminus (Christianson *et al.*, 2010). The NAC domain can be further subdivided into five subdomains [A-E], among which the subdomains C and D are highly conserved and may mainly act in DNA binding (Ooka *et al.*, 2003). In addition, the subdomain D of some NAC proteins contains a hydrophobic NAC repression domain (NARD), which can suppress the transcriptional activity of NAC TFs by suppressing their DNA-binding ability or nuclear localization ability. As a specific functional motif in NARD, LVFY can suppress the activity of both NAC TFs and other TFs by its hydrophobicity (Hao *et al.*, 2010).

NAC TFs are involved in many biological processes in plants such as organ development, secondary wall synthesis, and stress response. For instance, *NAC29* and *NAC31* regulate the downstream cellulose synthase (CESA) by activating the downstream TF *MYB61* to control the synthesis of secondary wall (Huang *et al.*, 2015). For response to stresses, especially abiotic stresses, the NAC TFs are dominant regulators. *RD26* gene is the first NAC TF identified as a regulator of abscisic acid (ABA) and jasmonate (JA) signaling during stress response in Arabidopsis (Fujita *et al.*, 2004). *SNAC1* is a stress-responsive NAC TF that confers drought tolerance to rice by closing stomata (Hu *et al.*, 2006), and *SNAC3* confers heat and drought tolerance through regulating reactive oxygen species (Fang *et al.*, 2015). *OsNAC2* was also reported to be involved in drought stress response mediated by ABA (Shen *et al.*, 2017b); besides, it plays divergent roles in different biological processes, such as regulating shoot branching (Mao *et al.*, 2007) and plant height and flowering time through mediating the gibberellic acid (GA) pathway (Chen *et al.*, 2015). *VNI2* is a bifunctional NAC TF reported as a transcriptional repressor of xylem vessel formation in Arabidopsis (Yamaguchi *et al.*, 2010), but the transcriptional activator activity of *VNI2* is induced under high salinity stress to regulate the leaf longevity (Yang *et al.*, 2011).

Nine seed-specific NAC TFs have been found through transcriptome analysis in rice (Mathew *et al.*, 2016). Here, four of these genes, *ONAC025, ONAC127, ONAC128* and *ONAC129*, which are located in the linked region of chromosome 11 and identified as a gene cluster, were selected for further analysis (Fang *et al.*, 2008). A recent study implied that two NAC TFs specifically expressed in maize seeds, *ZmNAC128* and *ZmNAC130*, play critical roles in starch and protein accumulation during grain filling (Zhang *et al.*, 2019). We therefore inferred that the four NAC TFs might also be involved in some important processes in rice seed development. To be more specific, we selected *ONAC127* and *ONAC129* from the four genes for study, which could form a heterodimer and participate in apoplasmic transport and heat stress response to regulate rice grain filling. The findings are expected to improve the understanding of the regulatory network of stress response and grain filling in rice.

## Materials and methods

### Generation of transgenic plants and growth conditions

For the generation of CRISPR mutants, the web-based tool CRISPR-P v2.0 (http://cbi.hzau.edu.cn/CRISPR2/) (Liu *et al.*, 2017) was used to obtain the specific gRNA cassettes targeting *ONAC127* and *ONAC129*, which were then cloned into a binary vector pYLCRISPR/Cas9-MH (Ma *et al.*, 2015). For overexpression plants, the stop-code-less cDNA fragments of *ONAC127* and *ONAC129* were amplified, and the 3×Flag or EGFP coding sequence was fused at the 3’ end. The fused sequences were cloned into the binary vector pCAMBIA1301U (driven by a maize ubiquitin promoter). For tissue specific expression analysis, a 2000-bp fragment of the 5’ upstream region of *ONAC127* and a 2043-bp fragment of the 5’ upstream region of *ONAC129* were cloned into the binary vector pDX2181G (with GUS, β-glucuronidase). These recombinant constructs were introduced into rice Zhonghua11 (ZH11; *Oryza sativa* ssp. *japonica*) by *Agrobacterium tumefaciens* (EH105)-mediated transformation (Lin and Zhang, 2005). The plants were cultivated in the paddy fields of Huazhong Agricultural University, Wuhan, China, under natural long-day conditions (approximately 12–14 h light/10–12 h dark) during May to October 2018. The temperature data of the growing area during rice reproductive stage are shown in Supplementary Fig. S7. The temperature of 35°C was set as the heat damage temperature of rice (T_B_), and the heat damage accumulated temperature per hour (TH_i_) was calculated as 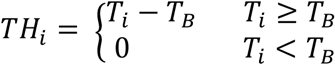 (T_i_ is the ambient temperature at *i* hour); heat damage hours during filling stage (H_S_) was calculated as 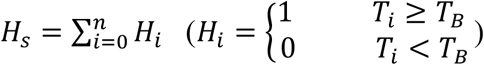; and heat damage accumulated temperature during filling stage (T_S_) was calculated as 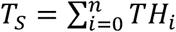 using the method described by Chen et al. (Chen *et al.*, 2019). The Ts during 0–7 DAP (T_S7_) and Hs during 0–7 DAP (H_S7_) were also calculated as the method above. Since two batches of rice plants were flowering in a large scale on about July 20^th^ and August 20^th^, we marked most of the flowering spikelets in all plants at the days around these two dates (including mutants, overexpression lines and ZH11(WT)), and the immature seeds used in the experiment were all from the spikelets marked in these days. Therefore, these two dates were set as the starting date for the calculation of the heat accumulation temperature. The phenotypes were detected in homozygous T1 generation of transgenic plants. The primer sequences used in this study are listed in Supplementary Table S1 and would not be repeated hereafter.

### Histochemical GUS staining

Immature seeds on 5 and 7 DAP from pONAC127::GUS and pONAC129::GUS transgenic plants were collected for GUS staining assay following the previously described method (Yang *et al.*, 2018).

### Microscopy analysis

For semi-thin section microscopy analysis, immature seeds on 7 DAP were placed in 2.5% glutaraldehyde for fixation, vacuum infiltrated on ice for 30 min and incubated at 4^°^C for 24 h. The plastic embedding and sectioning were performed as previously described (Wang *et al.*, 2008a). The slides were stained with toluidine blue and detected by a BX53 microscope (Olympus).

### RNA in situ hybridization

Immature seeds (1–10 DAP) from rice ZH11 were collected for paraffin embedding. Paraffin sectioning was performed according to a previous method (Xiong *et al.*, 2019). Gene-specific fragments of *ONAC127* and *ONAC129* were amplified by the primers used in RT-PCR and cloned into the pGM-T vector. The probes were synthesized using the DIG RNA labeling kit (SP6/T7) (Roche) according to the manufacturer’s recommendations. RNA hybridization and immunologic detection of the hybridized probes were performed on sections as described previously (Kouchi and Hata, 1993). Slides were observed using a BX53 microscope (Olympus).

### Transient expression assays

To investigate the subcellular localization of ONAC127 and ONAC129, the CDS of these genes was cloned into the pM999-35S vector, and the nuclear located gene *Ghd7* was used as a nuclear localization marker (Xue *et al.*, 2008). For the BiFC assay, the CDS of *ONAC127* and *ONAC129* was cloned into the vectors pVYNE and pVYCE (Waadt *et al.*, 2008), respectively. The plasmids were extracted and purified using the Plasmid Midi Kit (QIAGEN) and then transformed into rice protoplasts according to the previously described procedure (Shen *et al.*, 2017a). The fluorescent signals were detected with a confocal laser scanning microscope (TCS SP8, Leica).

### RNA isolation and transcript analysis

Total RNA was isolated using the TRIzol method (Invitrogen), and the first strand cDNA was synthesized using HiScript II Reverse Transcriptase (Vazyme). The real-time PCR was performed on QuantStudio 7 Flex Real-Time PCR System (Applied Biosystems), with the 2^ΔΔCt^ method for relative quantification (Livak and Schmittgen, 2001). The significance of differences was estimated using Student’s t-test. Relevant primers were designed according to qPrimerDB (https://biodb.swu.edu.cn/qprimerdb) (Lu *et al.*, 2018).

### Yeast assays

For yeast two-hybrid assay, the CDS of *ONAC127* and *ONAC129* was cloned into the pGBKT7 and pGADT7 vector (Clontech). The fusion plasmids were then transformed into yeast strain AH109 or Y187. The pGBKT7-53 was co-transformed with pGADT7-T as a positive control. For yeast one-hybrid assay, DNA fragments of about 1000 bp corresponding to the promoters of the target genes were separately inserted into the pHisi-1 plasmid (Clontech). *ONAC127* and *ONAC129* were fused to GAL4 transcriptional activation domain (pGAD424). These constructs were then transformed into the yeast strain YM4271. Yeast two-hybrid assay and one-hybrid assay were performed following the manufacturer’s instructions (Clontech).

### Chromatin immunoprecipitation (ChIP)

*ONAC127* and *ONAC129* overexpression lines (fused with 3×FLAG) were used for ChIP-Seq analysis. The expression of the target proteins was verified by western blot using ANTI-FLAG M2 Monoclonal Antibody (Sigma-Aldrich). ChIP assay was performed with ANTI-FLAG M2 Magnetic Beads (Sigma-Aldrich) with the previously described method (Xiong *et al.*, 2019). For each library, three independent replicated samples were prepared. The immunoprecipitated DNA and input DNA were subjected to sequencing on the HiSeq 2000 platform (Illumina) by the Novogene Corporation. The quality of the sequencing data is shown in Supplementary Dataset S3.

ChIP-Seq raw sequencing data were mapped to the rice reference genome (RGAP ver. 7.0, http://rice.plantbiology.msu.edu) (Kawahara *et al.*, 2013) using BWA (Li and Durbin, 2009). MACS2 (Zhang *et al.*, 2008) was used for peak calling and the peaks were identified as significantly enriched with corrected *p*-value < 0.05 in the IP libraries compared with input DNA. Visual analysis was performed using IGV (Intergative Genomics Viewer, v2.3.26) (Robinson *et al.*, 2011). Motif enrichment analysis was performed by MEME (Machanick and Bailey, 2011) with default parameters.

To validate the specific target genes, the immunoprecipitated DNA and input DNA were applied for ChIP-qPCR analysis. The enrichment value was normalized to that of the input sample. The significance of differences was estimated using Student’s t-test.

### RNA-Seq

Total RNA was extracted from immature seeds at 7 DAP. For each library, three independent replicate RNA samples were prepared. The RNA samples were then sequenced on the HiSeq 2000 platform (Illumina) by the Novogene Corporation. The quality of the sequencing data is shown in Supplementary Dataset S3.

The raw reads were filtered to remove the adaptors and low-quality reads. Clean reads were then mapped to the reference genome of rice (RGAP v. 7.0) using HISAT2 (v.2.0.5) (Kim *et al.*, 2015). The differentially expressed genes (DEGs) (ļFold Changeļ ≥ 2 and corrected *p*-value < 0.05) were selected by DESeq2 software (Love *et al.*, 2014). GOseq (Young *et al.*, 2010) was used for GO enrichment analysis, and FDR was converted to −log_10_(FDR) for display.

### In vitro GST pull-down assay

The CDS of *ONAC127* and *ONAC129* was cloned into pET28a and pGEX-4-1 vectors respectively for His-tagged ONAC127 and GST-tagged ONAC129 protein expression *in vitro*. The fusion plasmids were transformed into *Escherichia coli* BL21 strain. The GST pull-down assay was performed according to the previous method (Xiong *et al.*, 2019). The protein was separated on a 10% SDS-PAGE gel and further analyzed by immunoblotting using anti-His and anti-GST antibody (Sigma-Aldrich).

### Dual luciferase transcriptional activity assay

Full-length cDNAs of *ONAC127* and *ONAC129* were cloned to yeast GAL4 binding domain vectors (GAL4BD) and “None” as effectors. The 35S-GAL4-fLUC and 190-fLUC were used as reporters, and AtUbi::rLUC was used as an internal control. The constructed plasmids were purified and transformed into rice protoplasts with the procedure described above. Dual Luciferase Reporter Assay System (Promega) was used to measure the luciferase activity by Tecan Infinite M200 (Tecan).

### Agronomic trait analysis

The harvested mature rice grains were air dried and stored at room temperature for at least 2 months before measuring. Only full grains were used for measuring the 1000-grain weight, which was calculated based on 100 grains. The seed set rate/composition was calculated based on the grains from 3–5 panicles (including the full, shrunken and blighted grains). All measurements of the positive transgenic plants were performed with three independent lines.

### Accession Numbers

The sequence data in this study can be found in the RGAP database (http://rice.plantbiology.msu.edu) under the following accession numbers: *ONAC025* (*LOC_Os11g31330*), *ONAC127* (*LOC_Os11g31340*), *ONAC128* (*LOC_Os11g31360*), *ONAC129* (*LOC_Os11g31380*), *OsMST6* (*LOC_Os07g37320*), *OsSWEET4* (*LOC_Os02g19820*), *OsEATB* (*LOC_Os09g28440*), *OsMSR2* (*LOC_Os01g72530*), *bHLH144* (*LOC_Os04g35010*), *OsHCI1* (*LOC_Os10g30850*), *HSP101* (*LOC_Os05g44340*), *OsSWEET11* (*LOC_Os08g42350*), *OsSWEET15* (*LOC_Os02g30910*), *OsbZIP63* (*LOC_Os07g48820*). The RNA-seq and ChIP-seq data are deposited in the NCBI Gene Expression Omnibus (Edgar *et al.*, 2002) with the accession number of GSE140167.

## Results

### ONAC127 and ONAC129 were specifically expressed in rice seeds

For a basic understanding of *ONAC025, ONAC127, ONAC128* and *ONAC129*, we firstly checked the expression data of these four genes in the microarray database CREP (http://crep.ncpgr.cn) (Wang *et al.*, 2010). The results showed that *ONAC025* had significantly higher expression levels in various tissues than other genes, while *ONAC128* had the lowest expression level. It is noteworthy that the expression level of *ONAC127* and *ONAC129* in the seed reached the maximum at 7 day after pollination (DAP), followed by gradual decreases.

To validate the spatial and temporal expression patterns and further explore the specific distribution of *ONAC127* and *ONAC129* expression in rice seeds, we mechanically isolated rice seeds at 5–14 DAP into starch endosperm, embryo and the mixture of pericarp and aleurone layer for qRT-PCR analysis according to a previously described method (Bai *et al.*, 2016). The gene specifically expressed in the aleurone layer *oleosin* and that specifically expressed in starch endosperm *SDBE* were used as markers (Ishimaru *et al.*, 2015) (Supplementary Fig. S1). The results showed that *ONAC127* and *ONAC129* were predominantly expressed in the pericarp or aleurone layer, while weakly expressed in starch endosperm, and the expression reached the maximum level at 5 DAP during the whole development process (Fig. 1A).

**Fig. 1.**
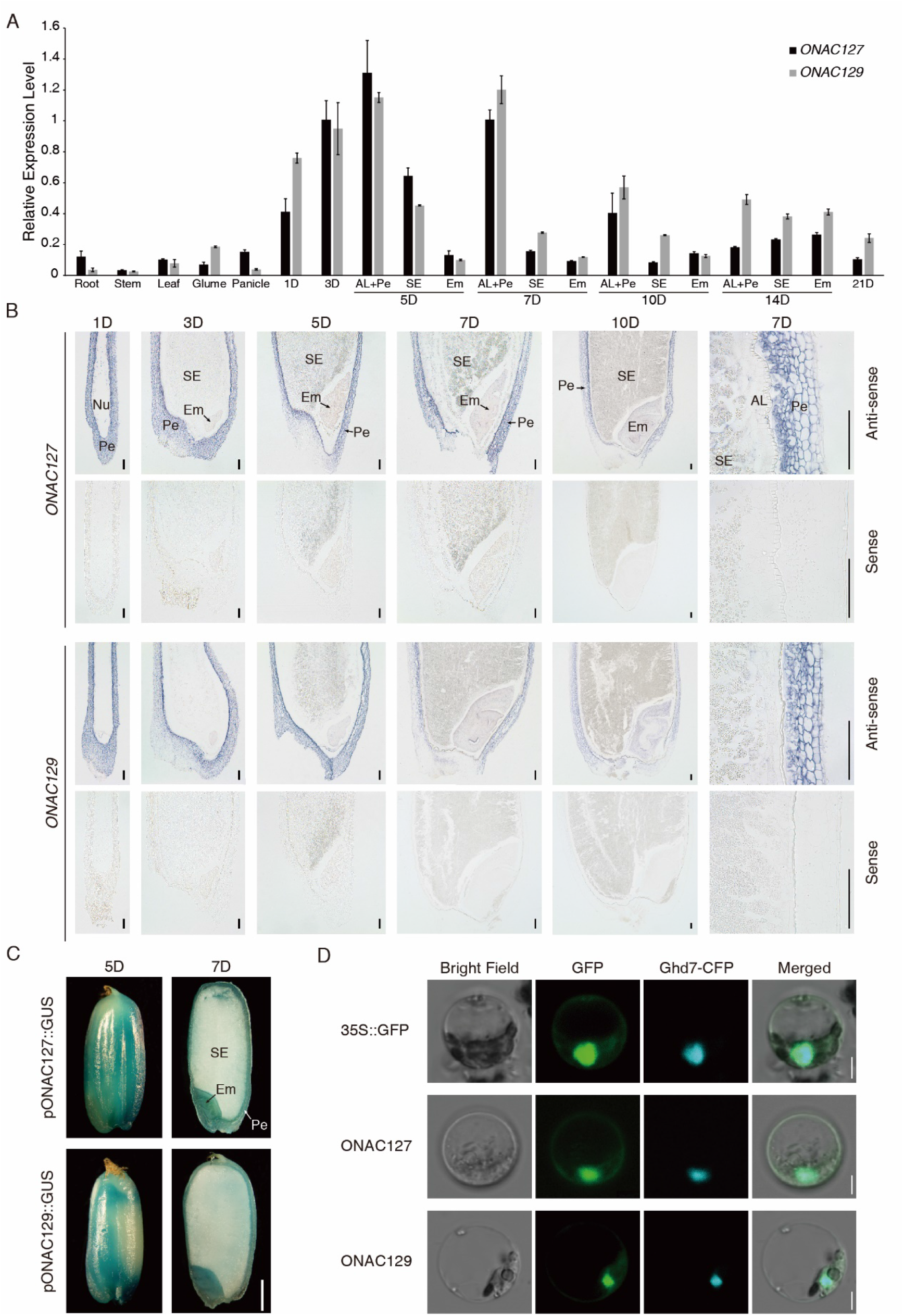
Spatial and temporal expression patterns of *ONAC127* and *ONAC129*. (A) qRT-PCR analysis of *ONAC127* and *ONAC129* in different tissues. AL, aleurone layer; Pe, pericarp; SE, starchy endosperm; Em, embryo; 1-21D, caryopses collected during 1–21 DAP. The values in each column are the mean of three technical replicates and the error bars indicate the ±SD. Ubiquitin was used as the reference gene. (B) *In situ* hybridization of sectioned seeds collected at 1, 3, 5, 7 and 10 DAP using anti-sense and sense probes. Nu, nucleus; Pe, pericarp; SE, starchy endosperm; Em, embryo; AL, aleurone layer. Scale bars = 50 μm. (C) Histochemical GUS activity detection using pONAC127::GUS and pONAC129::GUS. Pe, pericarp; SE, starchy endosperm; Em, embryo. Scale bar = 1 mm. (D) Subcellular localization of *ONAC127* and *ONAC129* in rice protoplasts. 35S::Ghd7-CFP was used as a nuclear marker. Scale bars = 5 μm.

The mRNA *in situ* hybridization analysis on the immature seeds of ZH11 showed that *ONAC127* and *ONAC129* were dominantly expressed in the pericarp, while their expression levels in the aleurone layer and starch endosperm were rather low (Fig. 1B). Histochemical GUS activity was also detected in the transgenic plants expressing pONAC127::GUS and pONAC129::GUS, and the results were almost identical to those of *in situ* hybridization analysis: *ONAC127* and *ONAC129* were predominantly expressed in the pericarp of seeds (Fig. 1C). Since 5 DAP is a key time point of seed development when the organogenesis is completed and the maturation stage just initiates (Agarwal *et al.*, 2011), *ONAC127* and *ONAC129* might be involved in some substance accumulation processes and play some vital roles in grain filling and maturation.

We then transiently transformed ONAC127 or ONAC129 protein with the fusion of green fluorescent protein (GFP) into rice protoplast to investigate the subcellular localization pattern of the two genes. 35S::GFP was used as a positive control and 35S::Ghd7-CFP was used as a nuclear marker (Xue *et al.*, 2008). The fluorescent signals generated by ONAC127-GFP and ONAC129-GFP were distributed in the nucleus and cytoplasm just like the positive control (Fig. 1D). To further confirm the subcellular localization, transgenic plants expressing pUbi::ONAC127-GFP and pUbi::ONAC129-GFP were generated in Zhonghua11 (ZH11) background. The diverse cell types and large number of starch grains in seeds made it difficult to distinguish the protein subcellular localization in rice seeds. Hence, the subcellular localization patterns of ONAC127 and ONAC129 were validated by detecting the fluorescent signals from the roots of two-week-old rice seedlings. The results showed that ONAC127 and ONAC129 proteins were indeed localized in the nucleus and cytoplasm (Supplementary Fig. S2).

### ONAC127 and ONAC129 formed a heterodimer

It has been reported that the NAC TFs usually function as a dimer (Olsen *et al.*, 2005). Thus, we investigated whether there is any interaction between ONAC127 and ONAC129. The results indicated that ONAC127 could interact with ONAC129 in yeast (Fig. 2A). We also determined whether ONAC127 and ONAC129 could form homodimers, but the results showed that they could not interact with themselves in yeast. To confirm the interaction between ONAC127 and ONAC129, *in vitro* GST pull-down assay was carried out. His-tagged ONAC127 and GST-tagged ONAC129 were expressed respectively and then incubated together. The protein mixture of GST and ONAC127-His was used as a negative control. After purification by Glutathione Agarose, ONAC127-His was detected in the sample containing ONAC129-GST instead of the control with His-tag antibody (Fig. 2B). *In vivo* bimolecular fluorescence complementation (BiFC) assay was also performed with rice protoplasts, with nuclear homodimer OsbZIP63 being chosen as the positive control (Walter *et al.*, 2004), and 35S::Ghd7-CFP being used as the nuclear marker. Yellow fluorescence generated from the interaction between ONAC127-YFP^N^ and ONAC129-YFP^C^ was detected, confirming the formation of a heterodimer by ONAC127 and ONAC129 in the nucleus (Fig. 2C). Previous studies have shown that some NAC TFs were located in the cytoplasm or plasma membrane, but under some specific conditions they could be imported into the nucleus (Fang *et al.*, 2014). Based on the subcellular localization patterns of ONAC127 and ONAC129 (Fig. 1D), we speculated that these two proteins might also exert their transcriptional regulation functions in this way.

**Fig. 2.**
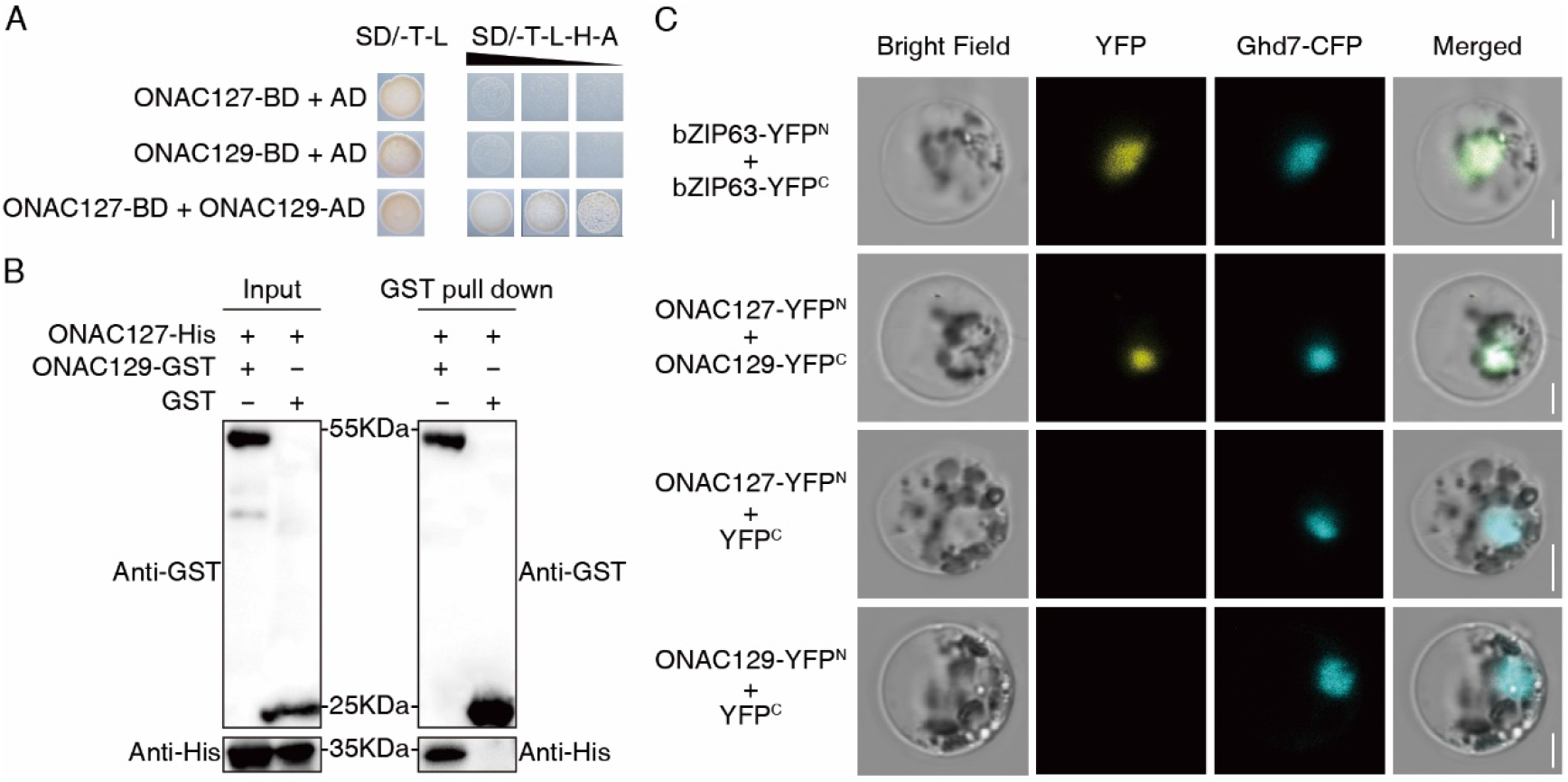
Interaction between ONAC127 and ONAC129. (A) Yeast two-hybrid assay. The full-length cDNA of ONAC127 and ONAC129 was both cloned into pGBKT7 (BD) and pGADT7 (AD). The transformants were grown on [SD/–Trp /–Leu] and [SD/–Trp/–Leu/–His/–Ade] plates with 10^0^, 10^-1^ and 10^-2^ fold dilution. (B) The pull-down assays showing that there is a direct interaction between ONAC127-His and ONAC129-GST *in vitro*. (C) The BiFC assays of ONAC127 and ONAC129. ONAC127-YFP^N^ and ONAC129-YFPC interact to form a functional YFP in rice protoplasts. 35S::Ghd7-CFP was used as a nuclear marker. Scale bars = 5 μm.

### ONAC127 and ONAC129 played key roles in starch accumulation during rice grain filling

*ONAC127* or *ONAC129* knockout mutants (*onac127* and *onac129*) were obtained in ZH11 background by CRISPR/Cas9 genome editing system (Ma *et al.*, 2015). Considering that these two proteins could form a heterodimer, double knockout mutants (*onac127;129*) were also obtained. The sgRNA target sites were designed at the exons of *ONAC127* and *ONAC129* by using the web-based tool CRISPR-P (Liu *et al.*, 2017). There were two target sites of each target gene in corresponding mutants, which were expected to generate different mutations in the coding region of corresponding target genes (Fig. 3A). The seeds of three T0 homozygous plants of each mutant were selected for the generation of independent T1 transgenic lines. The sequencing data of the transgenic lines were decoded by the method described in a previous study (Liu *et al.*, 2015), and the genotypes of the homozygous lines are shown in Supplementary Fig. S3. On the other hand, the overexpression lines pUbi::ONAC127-FLAG and pUbi::ONAC129-FLAG (OX127 and OX129) were also generated (Fig. 3B). The relative expression levels of *ONAC127* and *ONAC129* in the overexpression lines were analyzed by qRT-PCR (Supplementary Fig. S4).

**Fig. 3.**
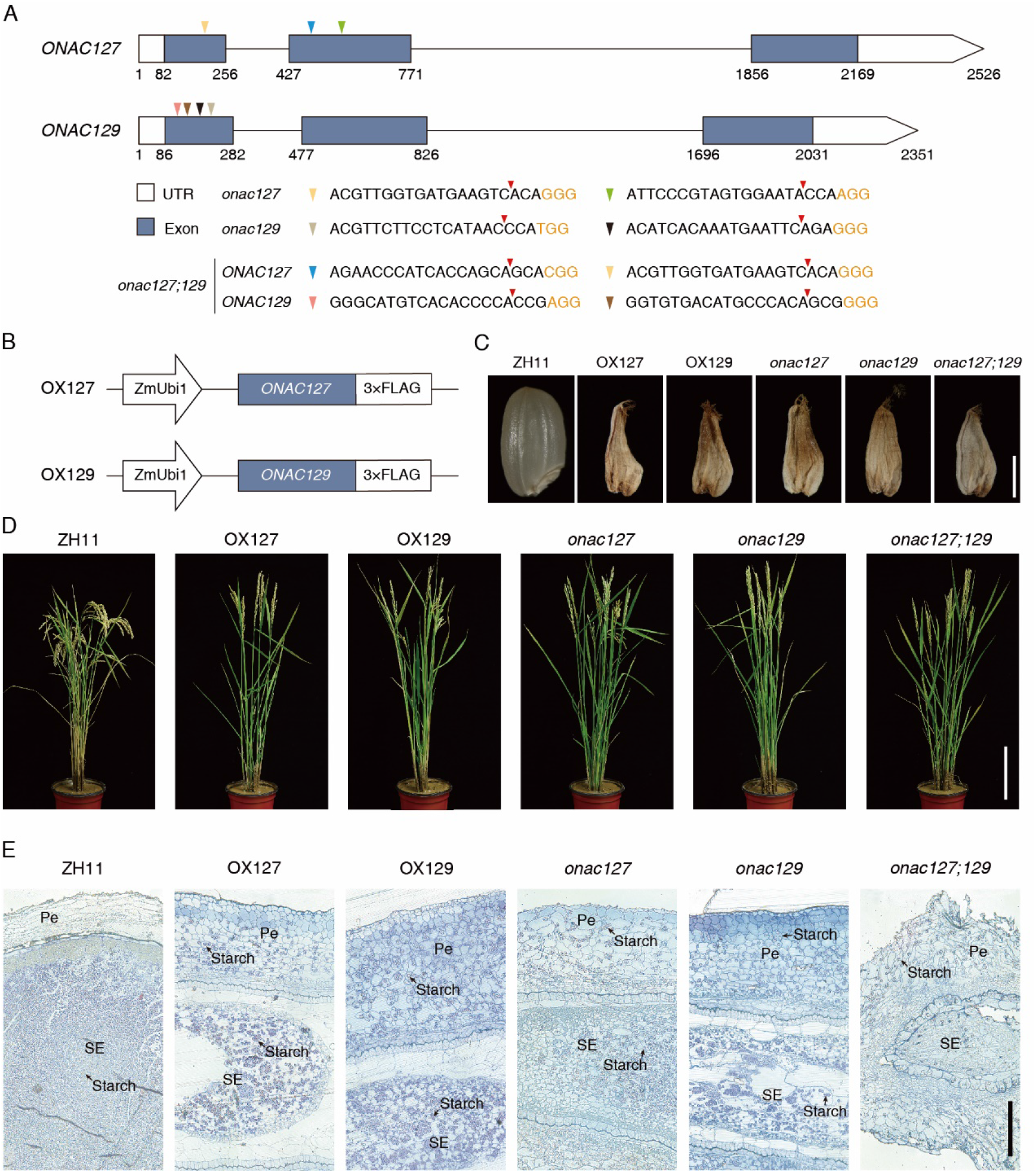
Construction of CRISPR mutants and overexpression plants and their phenotypes. (A) Schematic diagram of the sgRNA target sites in genomic region of *ONAC127* and *ONAC129*. The target sequences of *onac127, onac129* and *onac127;129* are listed under the diagram and marked as a colorful triangle. The Cas9 splice sites are marked as red triangle and the protospacer-adjacent motifs are shown in orange. (B) Schematic diagram of the construction of OX127 and OX129. (C) Phenotypes of grains of the overexpression lines and mutants of *ONAC127* and *ONAC129*. Scale bars = 2 mm. (D) Phenotypes of panicles at maturation stage. The panicles of transgenic plants are almost upright while those of ZH11 are bent due to grain filling. Scale bars = 20 cm. (E) Accumulation of starch in the seed pericarp of overexpression lines and mutants relative to ZH11. Cross sections of seeds stained with toluidine blue at 7DAP. Pe, pericarp; SE, starchy endosperm. Scale bar = 40 μm.

During the rice reproductive stage, it was found that the development of some seeds in both the overexpression lines and mutants of *ONAC127* and *ONAC129* was arrested from 5 DAP to 7 DAP (Supplementary Fig. S5), resulting in some shrunken and incompletely filled grains after maturation (Fig. 3C, D). As mentioned before, the expression level of *ONAC127* and *ONAC129* was very high during 5–7 DAP (Fig. 1A), and their functions seemed to be closely related to grain filling, especially at the early stage of maturation. Since a contradictory phenotype was observed, we determined the expression levels of *ONAC127* and *ONAC129* in the transgenic lines. As a result, *ONAC127* and *ONAC129* were up-regulated in the overexpression lines and down-regulated in the mutants (Supplementary Fig. S6). It can be speculated that ONAC127 and ONAC129 may regulate several different pathways to contribute to rice grain filling at the same time. The expression of ONAC127 and ONAC129 may need to be maintained at a steady and balanced level, and any fluctuation in their expression may interfere with the normal function of different pathways to cause defective grain filling in rice.

To further explore the reason for the phenotype of shrunken grains, histological analysis with resin embedded sections was carried out with 7 DAP seeds of incomplete filling. In ZH11, starch is transiently stored in the pericarp during the early stage, and starch degradation in the pericarp is correlated with starch accumulation in the endosperm (Wu *et al.*, 2016). As expected, neatly and densely arranged starch grains were found in the endosperm of ZH11 seeds, while hardly observed in the pericarp (Fig. 3E). In contrast, many starch grains were accumulated in the pericarp of the mutants and the overexpression lines, particularly in the double mutant *onac127;129*, in which few starch grains could be found in the endosperm (Fig. 3E). These findings suggested that *ONAC127* and *ONAC129* might play critical roles in the translocation and mobilization of starch to developing endosperm.

### ONAC127 and ONAC129 were involved in heat stress response

In the measurement of agronomic traits, the transgenic lines of *ONAC127* and *ONAC129* showed significant reduction in both seed setting rate (percentage of fully filled grains per panicle after harvest) and 1000-grain weight (only fully filled grains were weighed) compared with ZH11 (Table 1). The transgenic lines showed an obvious increase in the percentage of shrunken grains per panicle while no significant change in the proportion of blighted grains per panicle (glumes with no seed) (Table 1), proving that the reduction of seed setting rate and grain yield could be mainly ascribed to the increase in shrunken grains.

**Table 1.**
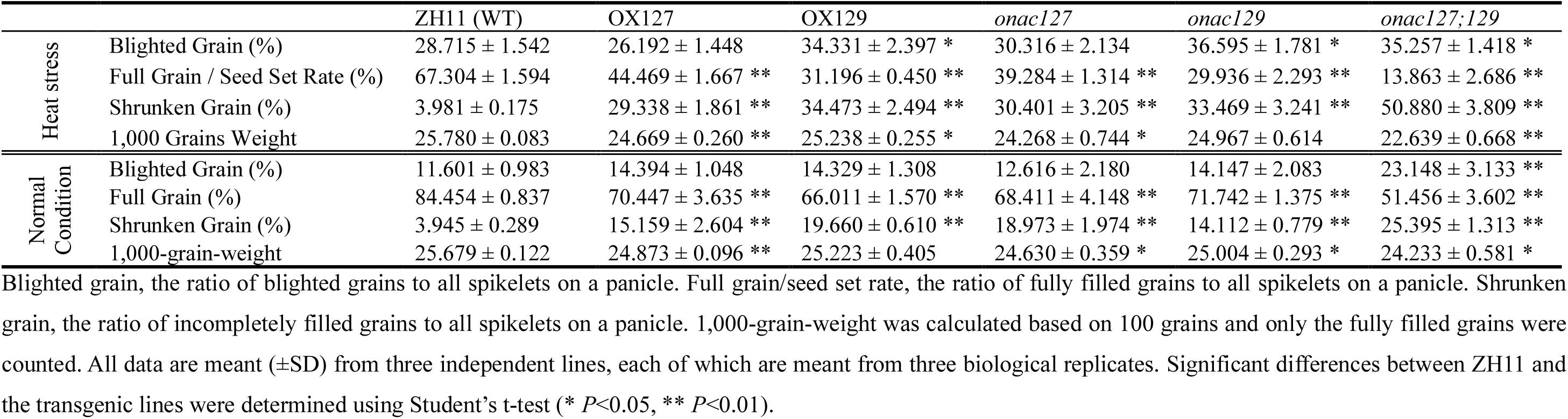
Agronomic trait analysis of wild type (ZH11) and transgenic lines of *ONACI27!ONACI29*.

Notably, the transgenic plants suffered from severe heat stress at the grain filling stage in the summer of 2018 (Supplementary Fig. S7). Previous studies have shown that the ambient temperature higher than 35°C at flowering and grain filling stages has a serious impact on the yield (Hakata *et al.*, 2017). Therefore, we set 35°C as the heat damage temperature of rice (T_B_), and calculated the heat damage accumulated temperature per hour (TH_i_), heat damage accumulated temperature during filling stage (T_S_) and heat damage hours during filling stage (H_S_) as previously described (Chen *et al.*, 2019). Given that *ONAC127* and *ONAC129* function at the early grain filling stage, we also calculated the heat damage accumulated temperature during 0–7 DAP (T_S7_) and heat damage hours during 0–7 DAP (H_S7_). It is worth noting that two batches of rice were cultivated in the summer of 2018, and all the rice plants in the same batch were sown on the same day. The first batch of plants flowered on about July 20^th^ while the second batch flowered on about August 20^th^. It is obvious that the T_S_ and H_S_ of the first batch were much higher than those of the second batch, especially at the early development stage of seeds (Supplementary Fig. S7). Therefore, the plants cultivated in the first batch were defined as suffering from heat stress, while those cultivated in the second batch were defined as growing under normal conditions with little or no heat stress. It was found that heat stress caused sharp increases of shrunken grains compared with normal conditions, and such increase was more dramatic in *onac127;129* than in other transgenic lines, which led to a significant decrease in the seed setting rate (Table 1). Meanwhile, the expression level of *ONAC127* and *ONAC129* was drastically increased under heat stress (Supplementary Fig. S8), suggesting that these two genes might be involved in heat stress response at the grain filling stage of rice.

### ONAC129 negatively regulated ONAC127 transcriptional activity

To further understand the relationship between *ONAC127* and *ONAC129*, it was necessary to determine the transcriptional activity of these two genes *in vivo*. Therefore, a dual luciferase assay was performed in rice protoplasts. *ONAC127* and *ONAC129* were respectively fused with the yeast GAL4 binding domain (GAL4BD) as effectors, which were then co-transformed into rice protoplasts with a firefly luciferase reporter gene driven by a CaMV35S promoter (35S-GAL4-fLUC; Fig. 4A). A significant increase was detected in the transcriptional activation activity of ONAC127 and ONAC129, while the transcriptional activity of ONAC127 was significantly suppressed when ONAC129 was co-transformed into rice protoplasts (Fig. 4B). These results suggested that ONAC129 might negatively regulate the transcriptional activity of ONAC127. We then attempted to determine whether ONAC129 and ONAC127 regulate the expression of each other. The results showed that there were no significant fluctuations in the expression of *ONAC127* in *ONAC129* transgenic lines and vice versa (Supplementary Fig. S6).

**Fig. 4.**
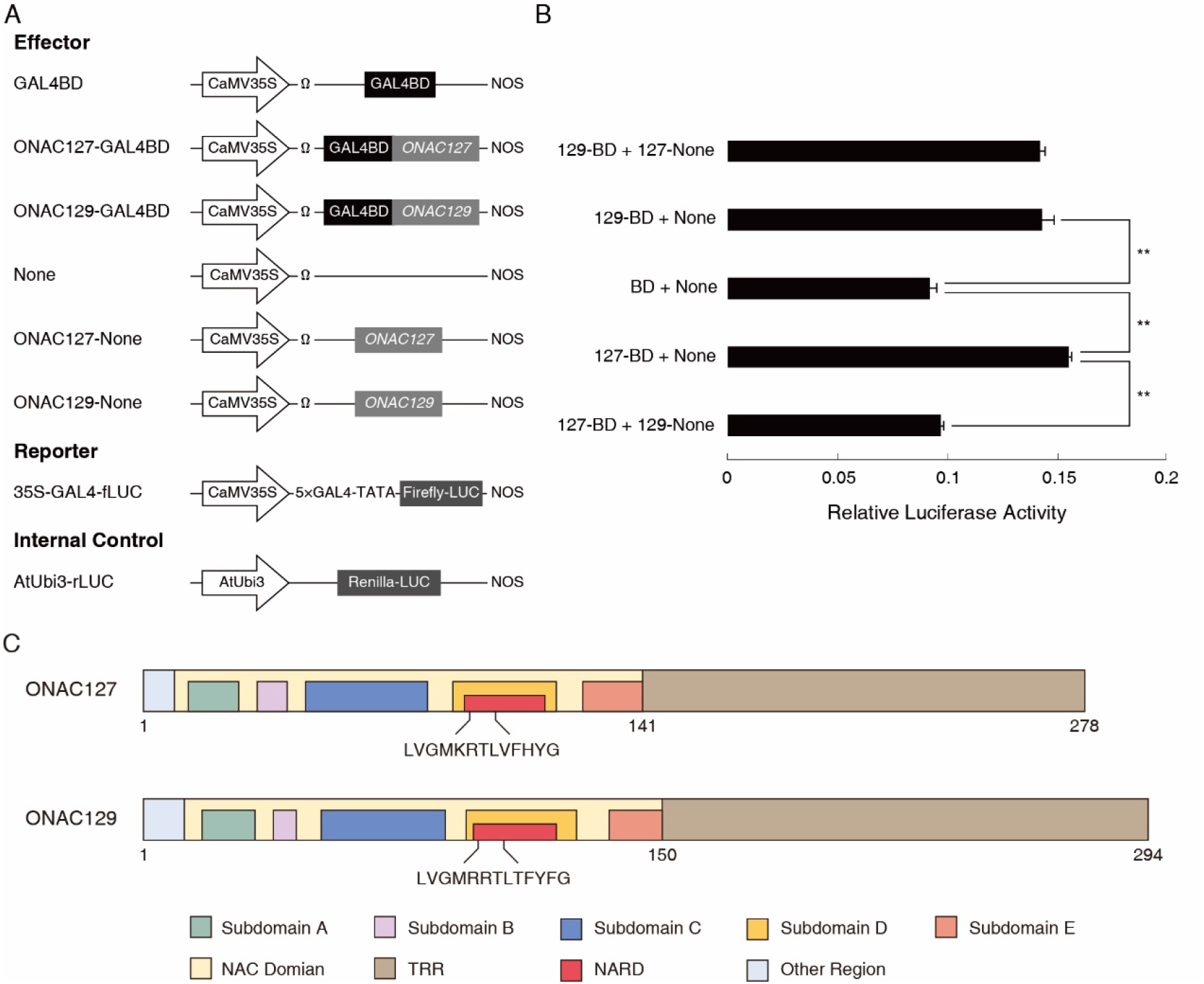
Negative regulation of ONAC129 on ONAC127 transcriptional activity. (A) Scheme of the constructs used in the rice protoplast cotransfection assay. (B) Data of Dual-LUC assay. The reporter and internal control were co-transformed with each experimental group. The fLUC/rLUC ratio represents the relative activity of CaMV35S promoter. The values in each column are the mean (±SD) of three replicates. Significant differences were determined using Student’s t-test (** *P*<0.01). (C) Schematic diagram of the domains of ONAC127 and ONAC129. The color legends indicating the domains shown at the bottom of the figure.

It is known that several NAC TFs contain both the transcriptional activation domain and NAC repression domain (NARD), which may determine the downstream events (Hao *et al.*, 2010). Hence, BLAST analysis was performed to compare the protein sequences of ONAC127 and ONAC129, and NARD-like sequences were found in the subdomain D of the NAC domain in both genes (Fig. 4C), suggesting that *ONAC127* and *ONAC129* might be bifunctional TFs that could activate or repress downstream genes in different situations.

### ONAC127 and ONAC129 regulated the transcription of genes related to sugar transportation and abiotic stimuli

RNA-Seq analysis can help to identify DEGs. RNA-seq libraries were generated with 7-DAP seeds of the overexpression lines and mutants of *ONAC127* and *ONAC129* under heat stress and normal conditions. Compared with ZH11 (ZH11H), a total of 7056, 6485, 5765, and 1608 DEGS were identified in *onac127 (onac127H), onac129 (onac129H*), OX127 (OX127H), and OX129 (OX129H) respectively under heat stress; compared with ZH11 (ZH11N), a total of 15334, 13284, 1776, and 2041 DEGs were identified in *onac127* (*onac127*N), *onac129* (*onac129*N), OX127 (OX127N), and OX129 (OX129N) respectively under normal conditions (Fig. 5A, B; Supplementary Dataset S1). Gene Ontology (GO) enrichment analysis was performed to examine the biological roles of these DEGs in seed development. As a result, the significantly enriched GO terms were mainly associated with stimulus response, transcriptional activity regulation, signal transduction, cell wall construction and substance transport (Fig. 5C).

**Fig. 5.**
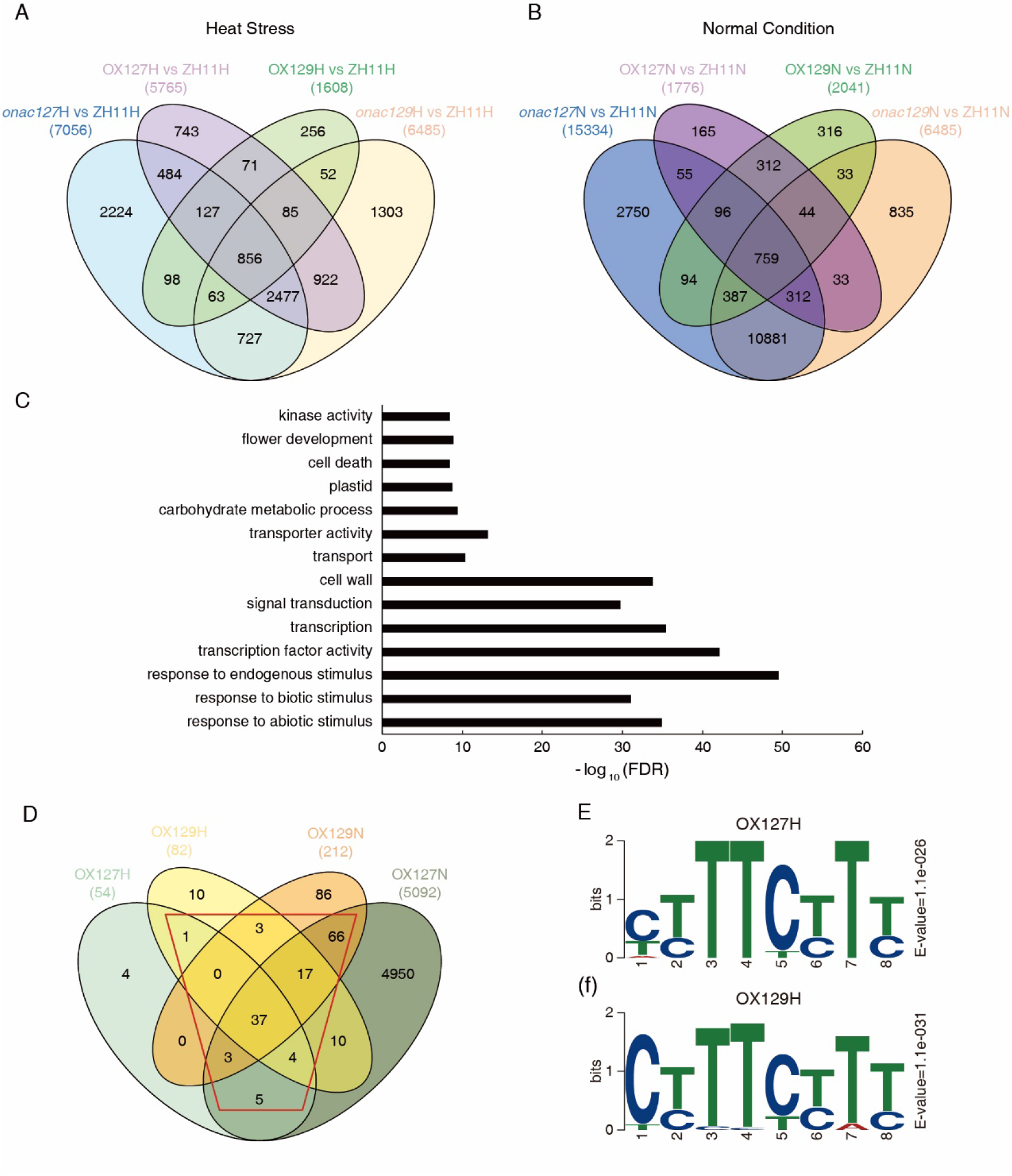
Data analysis of RNA-Seq and ChIP-Seq. (A) and (B): Venn diagram showing the number of genes regulated by *ONAC127* and *ONAC129* based on the RNA-seq analysis. Differentially expressed genes are defined as |Fold Change|≥2 and *p*-value < 0.05. *p*-values were calculated using the Fisher’s exact test. (C) A selection of enriched gene ontology (GO) terms of the differentially expressed genes. GO terms with FDR <0.05 were kept. The length of the bars represents the negative logarithm (base 10) of FDR. (D) Venn diagram showing the number of the direct target genes of ONAC127 and ONAC129. The red trapezoid indicates the preferably bound genes of ONAC127 and ONAC129. (E) and (F): Motif analysis of ONAC127 and ONAC129 binding peaks. The E-value is the enrichment *P*-value multiplied by the number of candidate motifs tested.

Subsequently, the genes directly bound by ONAC127 and ONAC129 in seeds were identified using Chromatin Immunoprecipitation Sequencing assays (ChIP-seq). The ChIP assays were performed using the anti-FLAG antibody with 7-DAP seeds of OX127 (OX127H) and OX129 (OX129H) under heat stress, and OX127 (OX127N) and OX129 (OX129N) under normal conditions. The expression of the ONAC127-FLAG and ONAC129-FLAG fusion proteins was verified by western blot analysis to validate the effectiveness of the FLAG tags (Supplementary Fig. S9). After sequencing, 185, 6125, 220 and 455 peaks were finally obtained respectively in OX127H, OX127N, OX129H and OX129N (Supplementary Dataset S2), and 54, 5092, 82 and 212 putative genes directly bound by ONAC127 and ONAC129 were identified in OX127H, OX127N, OX129H and OX129N respectively according to the peaks (Fig. 5D). Then, the binding motifs of ONAC127 and ONAC129 were identified using MEME (Machanick and Bailey, 2011), and the results showed that the significantly enriched motif was ‘CT(C)TTCT(C)TT’ (Fig. 5E, F), which was in line with ‘TT(A/C/G)CTT’, the specific motif of transmembrane NAC TFs *NTL6* and *NTL8* (Lindemose *et al.*, 2014). Both the two genes are associated with stimulus response, implying that *ONAC127* and *ONAC129* are probably involved in stress response. Notably, the motif ‘CT(C)TTCT(C)TT’ was only found to be significantly enriched in OX127H and OX129H, while there was no significantly enriched motif in OX127N and OX129N, implying that the target genes of ONAC127 and ONAC129 might be more specific under heat stress.

To further explore the target genes of ONAC127 and ONAC129, the genes consistently bound by ONAC127/129 or simultaneously bound by these two proteins under heat stress or normal conditions were defined as the genes preferably bound by ONAC127/129 (Fig. 5D, red trapezoid). Based on the data of RNA-Seq analysis and phenotype of transgenic lines of ONAC127 and ONAC129, eight genes associated with sugar transport or abiotic stress response were selected from the 136 preferably bound genes as the potential target genes (PTGs) of ONAC127 and ONAC129 for further analysis.

### ONAC127 and ONAC129 played pivotal roles in grain filling through directly regulating key factors in substance accumulation and stress response

Yeast one-hybrid assay was performed to validate the interactions between ONAC127/129 and the promoters of their PTGs. The results demonstrated that ONAC127 and ONAC129 could directly bind to the promoter sequences of *OsEATB, OsHCI1, OsMSR2, OsSWEET4, bHLH144* and *OsMST6* in yeast (Fig. 6A). Dual-LUC assay in rice protoplasts was performed to determine whether ONAC127 and ONAC129 directly affect the transcription of these genes. ONAC127 and ONAC129 were co-transformed into rice protoplasts with a firefly luciferase reporter gene driven by target gene promoters (Fig. 6B). It turned out that ONAC127 activated *OsMSR2* and *OsMST6* promoters significantly, and both ONAC127 and ONAC129 strongly repressed the *OsEATB* and *OsSWEET4* promoters *in vivo*. It is noteworthy that ONAC129 could suppress the transcriptional activation activity of ONAC127 on the promoters of *OsMSR2* and *OsMST6*, while could not regulate the transcription of *OsMSR2* and *OsMST6* (Fig. 6C).

**Fig. 6.**
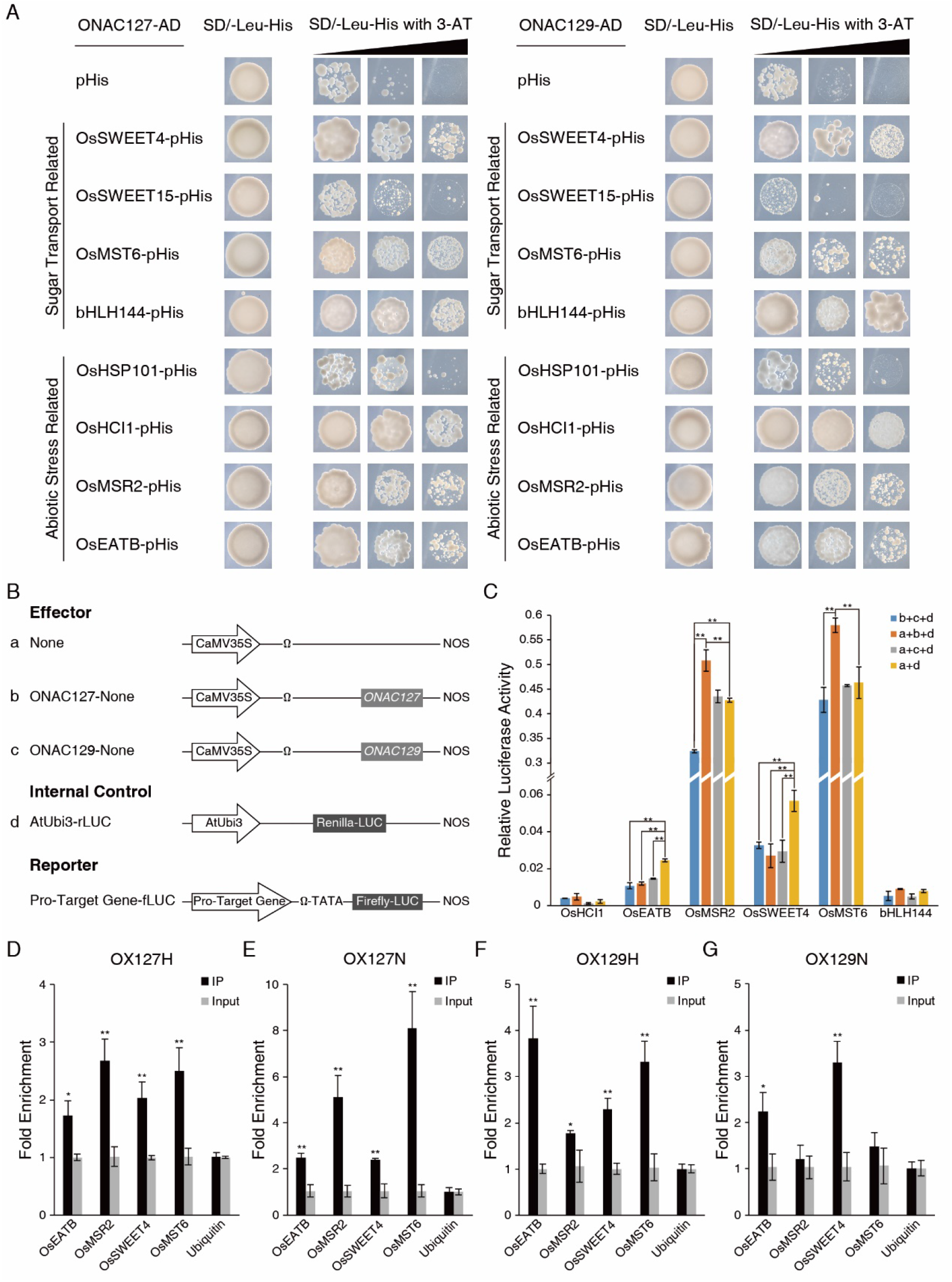
Identification and validation of the direct target genes of ONAC127 and ONAC129 in rice. (A) Interaction between ONAC127/129 and the promoters of PTGs as determined by yeast one-hybrid analysis. The transformants were grown on [SD/–Leu /–His] plates with 3-AT at concentrations of 50 mM, 100 mM and 150 mM, respectively. (B) Scheme of the constructs used in the rice protoplast cotransfection assay. (C) Data of Dual-LUC assay. The fLUC/rLUC ratio represents the relative activity of target gene promoters. (D-G) ChIP-qPCR verification of ONAC127 and ONAC129 binding regions. The values in each column are the mean (±SD) of three replicates. Significant differences were determined using Student’s t-test (* *P*<0.05; ** *P*<0.01).

To test whether the endogenous ONAC127 and ONAC129 can specifically bind their target genes, ChIP-qPCR was performed using the same rice materials of ChIP-Seq. The results indicated that ONAC127 and ONAC129 could bind to the promoters of *OsEATB*, *OsMSR2*, *OsSWEET4* and *OsMST6* under heat stress and normal conditions (Fig. 6D, E, F, G). The qRT-PCR for these genes was also performed in 7-DAP seeds of transgenic lines. The results revealed that the expression of *OsEATB* and *OsSWEET4* was significantly up-regulated in the mutants while generally down-regulated in the overexpression lines. The expression of both *OsMSR2* and *OsMST6* was up-regulated in *onac129* and down-regulated in *onac127* (Supplementary Fig. S10). These results indicated that *OsEATB*, *OsMSR2*, *OsSWEET4* and *OsMST6* are the direct targets of ONAC127 and ONAC129 in rice during grain filling.

## Discussion

In this study, two rice seed-specific NAC TFs *ONAC127* and *ONAC129* were identified, which can form a heterodimer and are predominantly expressed at the early and middle stage of rice seed development (Fig. 1, 2). Some grains of the transgenic plants were obviously unfilled (Fig. 3C, D, E; Supplementary Fig. S5), and there was a higher proportion of shrunken grains under natural heat stress (Table 1). Protein-DNA binding assays revealed that ONAC127 and ONAC129 may directly regulate the expression of the sugar transporters *OsMST6* and *OsSWEET4*. Both genes are involved in the transport of photosynthate from dorsal vascular bundles to endosperm during rice grain filling (Wang *et al.*, 2008b, Sosso *et al.*, 2015). ONAC127 and ONAC129 also bind to the promoters of *OsMSR2* and *OsEATB* directly (Fig. 6), which participate in abiotic stress response by responding to calcium ion (Ca^2+^) or plant hormones (Qi *et al.*, 2011, Xu *et al.*, 2011). These findings suggest that *ONAC127* and *ONAC129* may be involved in multiple pathways, and therefore interfere with the abiotic stress response and grain filling process at rice reproductive stage.

### ONAC127 and ONAC129 are involved in the apoplasmic transport of photosynthates as a balancer

At the early stage of seed development, some of the photosynthates from the dorsal vascular bundles are transported to the endosperm through the apoplasmic space formed by degenerative nucellar cell wall, and other photosynthates are used to synthesize starch grains in mesocarp cells (Hoshikawa, 1984). Starch accumulation in the pericarp will reach the maximum level at 5 DAP. After that, the starch in mesocarp cells is disintegrated and the nutrients are transported to the endosperm with the continuous filling of the endosperm cells. The inclusions of the pericarp cells will gradually disappear, and the cells will eventually dehydrate and die, only leaving cuticularized cell wall remnants (Wu et al., 2016).

Due to the abnormal expression of *ONAC127* and *ONAC129*, there may be no sufficient photosynthate to fill the endosperm cells in abnormal grains owing to the functional defect of the apoplasmic transport pathway. The starch grains formed by photosynthates accumulated in the pericarp are not disintegrated, which hinders the proliferation and filling of the endosperm cells by starch and eventually leads to incompletely filled and shrunken grains (Fig. 3C, E).

It is noteworthy that the target transporters *OsSWEET4* and *OsMST6* are predominantly expressed at the early stage of seed development, and may act downstream of the cell wall invertase OsCIN2 to import hexose into the aleurone layer through the apoplasmic space (Wang *et al.*, 2008b, Sosso *et al.*, 2015, Yang *et al.*, 2018). As mentioned above, ONAC127 and ONAC129 directly suppress the expression of *OsSWEET4* (Fig. 6C; Supplementary Fig. S10); besides, *ossweet4-1*, the mutant of *OsSWEET4*, showed a distinct phenotype of shrunken grains, which is similar to the abnormal grains in the transgenic lines of *ONAC127* and *ONAC129* (Sosso *et al.*, 2015). These facts confirm our hypothesis that *ONAC127* and *ONAC129* are involved in the apoplasmic transport pathways by regulating the expression of transporters.

Moreover, the shrunken grains in the mutants and overexpression lines of *ONAC127* and *ONAC129* are also similar to the incompletely filled grains of the double mutant *ossweet11;15*, in which two apoplasmic transporters OsSWEET11 and OsSWEET15 were dysfunctional (Yang *et al.*, 2018). Since *OsSWEET15* has been excluded from the direct target genes by yeast one-hybrid assays (Fig. 6A), we determined the expression of *OsSWEET11* and *OsSWEET15* in the transgenic lines of *ONAC127* and *ONAC129* by qRT-PCR. The results showed that both genes might be activated by ONAC127 and ONAC129 indirectly (Supplementary Fig. S11), suggesting that ONAC127 and ONAC129 might activate the expression of *OsSWEET11/15* while suppress that of *OsSWEET4* simultaneously. This may explain the result that the mutants and overexpression lines almost showed the same phenotype of incompletely filled grains (Fig. 3C, D, E; Supplementary Fig. S5). Hence, it can be speculated that *ONAC127* and *ONAC129* may act as a balancer in apoplasmic transport by activating or suppressing the expression of different transporters in a dynamic way to satisfy the demand of rice seed development at different stages (Fig. 7).

**Fig. 7.**
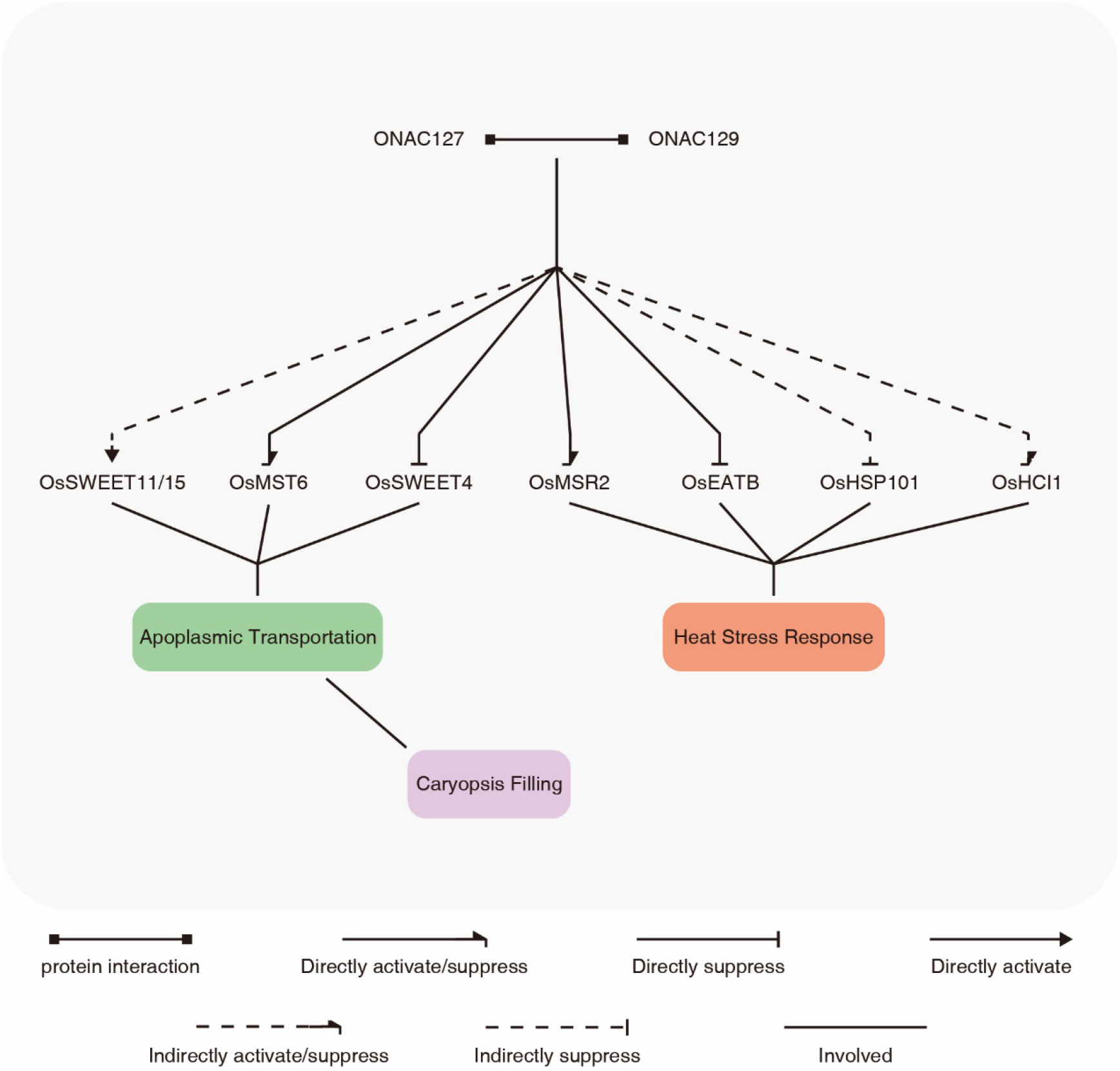
Schematic diagram of the working model of ONAC127 and ONAC129 in rice seeds. ONAC127 and ONAC129 directly regulate the expression of *OsMST6* and *OsSWEET4* to participate in the apoplasmic transport, and OsSWEET11 and OsSWEET15 may also be involved in this process as the indirect downstream transporters of them. When heat stress occurs, ONAC127 and ONAC129 may be involved in the stress response during the filling stage by regulating the expression of *OsMSR2* and *OsEATB* directly, while *OsHCI1* and *OsHSP101* are regulated indirectly. ONAC127 and ONAC129 may act as the balancer of apoplasmic transport and core transcription factors in the stress response during the early and middle stages of rice seed development.

ONAC129 suppresses the transcriptional activity of ONAC127 in some pathways. Besides, the regulation of individual ONAC127 and ONAC129 on downstream genes may be different from that of the heterodimer formed by them (Fig. 4C; Fig. 6C). We conjecture that the proportion of ONAC127/129 heterodimer may be vital to the balance of the amount of apoplasmic transporters at different stages of rice seed development. Whether it be knockout or overexpression of *ONAC127* or *ONAC129*, the proportion of the monomers and dimers of ONAC127/129 will be inevitably altered, which will induce various changes in the expression of the apoplasmic transporters and further cause the defects of apoplasmic transport, leading to the phenotype of incomplete grain filling (Fig. 3C). From a macroscopic point of view, the balanced regulation of apoplasmic transport may be a “rate-limiting step” of rice reproductive growth to ensure that there are sufficient photosynthates for both grain filling and plant energy to avoid premature aging, which is vital for the plants to keep a balance between vegetative growth and reproductive growth.

### ONAC127 and ONAC129 respond to heat stress during grain filling

Heat stress has profound effects on plant growth, particularly on the reproductive development. When plants encounter heat stress, the most rapid response is the rise of cytosolic Ca^2+^ concentration (Liu *et al.*, 2003). With the increase in Ca^2+^ concentration, the expression of calmodulins (CaMs) and calmodulin-like genes (CMLs) will be activated, and the phosphorylation state of HSFs will be modulated to regulate their DNA binding ability, which thereby regulates the expression of heat stress-related genes including HSPs and some plant hormone responsive genes (Li *et al.*, 2018).

The proportion of incompletely filled grains under heat stress was much higher than that under normal conditions (Table 1), and the expression of *ONAC127* and *ONAC129* was induced by heat stress (Supplementary Fig. S8), suggesting that heat stress might interfere with the dimerization of ONAC127 and ONAC129 to affect the balance of apoplastic transport. As mentioned above, ONAC127 and ONAC129 regulate the expression of *OsMSR2* and *OsEATB* directly (Fig. 6C; Supplementary Fig. S10). A previous study has shown that *OsMSR2*, which can be strongly induced by heat, cold and drought stress, is a calmodulin-like protein working by binding Ca^2+^ and acts as a Ca^2+^ sensor in plant cells (Xu *et al.*, 2011). *OsEATB* is a rice AP2/ERF gene, which can suppress gibberellic acid (GA) synthesis, while the GA signaling pathway is involved in the regulation of cell elongation and morphogenesis under heat stress (Koini *et al.*, 2009, Qi *et al.*, 2011). We hereby speculate that *ONAC127* and *ONAC129* might participate in heat stress response by regulating the Ca^2+^ and GA signaling pathways.

The expression level of *OsHSP101* and *OsHCI1* was significantly altered in the transgenic lines of *ONAC127* and *ONAC129* (Fig. 6A, C; Supplementary Fig. S11). Therefore, these two genes are also the potential target genes of ONAC127 and ONAC129, though they may not be regulated directly. RING E3 ligase gene *OsHCI1* is induced by heat stress and mediates nuclear-cytoplasmic trafficking of nuclear substrate proteins via mono-ubiquitination to improve the heat tolerance under heat shock (Lim *et al.*, 2013). Meanwhile, OsHSP101 functions as one of the important molecular chaperones that interact with OsHSA32 and OsHsfA2c, the latter of which is one of the central regulators of heat stress response (Singh *et al.*, 2012, Lin *et al.*, 2014). Accordingly, we speculate that *ONAC127* and *ONAC129* are involved in some core reactions in heat stress response at rice seed development stage.

Ca^2+^ is one of the most important second messengers in heat stress response, and cytosolic Ca^2+^ concentration will change rapidly when heat stress occurs (Li *et al.*, 2018). The stress signals will be then transmitted by CaMs/CMLs like OsMSR2 to influence the downstream gene expression. When the downstream HSFs are activated, HSPs including OsHSP101 will be induced, resulting in the occurrence of some nuclear-cytoplasmic trafficking reactions involving OsHCI1 (Singh *et al.*, 2012, Lim *et al.*, 2013). Besides, a variety of plant hormones including GA act in heat stress response to maintain plant growth, and *OsEATB* plays an important role in GA-mediated plant cell elongation (Qi *et al.*, 2011, Li *et al.*, 2018). Since all these direct or indirect target genes of *ONAC127* and *ONAC129* play vital roles in the key steps of heat stress response regulatory network, we speculate that ONAC127 and ONAC129 may be core TFs in heat stress response at rice graining filling stage, and the stress and hormone response pathways may be important in the complex regulatory network of apoplasmic transport.

## Abbreviations

DAP: Day after pollination
NAM, ATAF1/2, CUC2: NAC
NARD: NAC repression domain
PTGs: Potential target genes
TF: Transcription factor
ZH11: Zhonghua11

## Supplementary data

**Fig. S1.** Expression levels of the markers for the mechanical isolation of different tissues in rice seeds.

**Fig. S2.** Subcellular localization of ONAC127 and ONAC129 in the roots of two-week-old rice seedlings of pUbi::ONAC127-GFP and pUbi::ONAC129-GFP transgenic plants.

**Fig. S3.** Mutation sites in *onac127, onac129* and *onac127;129* lines as compared with wild-type (ZH11) sequences.

**Fig. S4.** Relative expression levels of *ONAC127* and *ONAC129* in overexpression lines.

**Fig. S5.** Different stages of seed development.

**Fig. S6.** Relative expression levels of *ONAC127* and *ONAC129* in 7-DAP seeds of transgenic lines compared with ZH11.

**Fig. S7.** Ambient temperature of the growing area at rice grain filling stage.

**Fig. S8.** Relative expression levels of *ONAC127* and *ONAC129* in 7-DAP seeds of ZH11 under heat stress and normal conditions.

**Fig. S9.** Detection of FLAG fusion proteins in ZH11 and overexpression lines.

**Fig. S10.** Expression levels of the target genes of ONAC127 and ONAC129 in 7-DAP seeds of transgenic lines compared with ZH11.

**Fig. S11.** Expression levels of the indirect target genes of ONAC127 and ONAC129 in 7-DAP seeds of transgenic lines compared with ZH11.

**Table S1.** Primers used in this study.

**Dataset S1.** Differentially expressed genes.

**Dataset S2.** Binding sites identified by ChIP-seq.

**Dataset S3.** Quality of Sequencing Data.

## Acknowledgements

We thank Prof. Min Chen, Prof. Honghong Hu, Prof. Yidan Ouyang and Ms. Wenjing Guo for helping revise the manuscript. This research was supported by grants from the National Natural Science Foundation of China (no. 31570321). The funders had no role in the study design, data collection and analysis, the decision to publish, or in the preparation of the manuscript.

## References

Agarwal, P., Kapoor, S. and Tyagi, A.K. (2011) Transcription factors regulating the progression of monocot and dicot seed development. Bioessays, 33, 189–202.

Bai, A.N., Lu, X.D., Li, D.Q., Liu, J.X. and Liu, C.M. (2016) NF-YB1-regulated expression of sucrose transporters in aleurone facilitates sugar loading to rice endosperm. Cell Res., 26, 384–388.

Chen, S., Zhang, T., Huang, Q., Chen, M., Zhao, L. and Zhang, Y. (2019) Effects of High Temperature Injury on Seed Setting Rate from Heading to Milky Stage of Early Season Rice in Hainan. China Tropical Agriculture, 65–69 (in Chinese with English abstract).

Chen, X., Lu, S., Wang, Y., Zhang, X., Lv, B., Luo, L., Xi, D., Shen, J., Ma, H. and Ming, F. (2015) OsNAC2 encoding a NAC transcription factor that affects plant height through mediating the gibberellic acid pathway in rice. Plant J., 82, 302–314.

Christianson, J.A., Dennis, E.S., Llewellyn, D.J. and Wilson, I.W. (2010) ATAF NAC transcription factors: regulators of plant stress signaling. Plant Signal Behav, 5, 428–432.

Edgar, R., Domrachev, M. and Lash, A.E. (2002) Gene Expression Omnibus: NCBI gene expression and hybridization array data repository. Nucleic Acids Res., 30, 207–210.

Fang, Y., Liao, K., Du, H., Xu, Y., Song, H., Li, X. and Xiong, L. (2015) A stress-responsive NAC transcription factor SNAC3 confers heat and drought tolerance through modulation of reactive oxygen species in rice. J. Exp. Bot., 66, 6803–6817.

Fang, Y., Xie, K. and Xiong, L. (2014) Conserved miR164-targeted NAC genes negatively regulate drought resistance in rice. J. Exp. Bot., 65, 2119–2135.

Fang, Y., You, J., Xie, K., Xie, W. and Xiong, L. (2008) Systematic sequence analysis and identification of tissue-specific or stress-responsive genes of NAC transcription factor family in rice. Mol. Genet. Genomics, 280, 547–563.

Fujita, M., Fujita, Y., Maruyama, K., Seki, M., Hiratsu, K., Ohme-Takagi, M., Tran, L.S., Yamaguchi-Shinozaki, K. and Shinozaki, K. (2004) A dehydration-induced NAC protein, RD26, is involved in a novel ABA-dependent stress-signaling pathway. Plant J., 39, 863–876.

Hakata, M., Wada, H., Masumoto-Kubo, C., Tanaka, R., Sato, H. and Morita, S. (2017) Development of a new heat tolerance assay system for rice spikelet sterility. Plant Methods, 13, 34.

Hao, Y.J., Song, Q.X., Chen, H.W., Zou, H.F., Wei, W., Kang, X.S., Ma, B., Zhang, W.K., Zhang, J.S. and Chen, S.Y. (2010) Plant NAC-type transcription factor proteins contain a NARD domain for repression of transcriptional activation. Planta, 232, 1033–1043.

Hoshikawa, K. (1984) Development of Endosperm Tissue with Special Reference to the Translocation of Reserve Substances in Cereals: III. Translocation pathways in rice endosperm. Japanese journal of crop science, 53, 153–162.

Hu, H., Dai, M., Yao, J., Xiao, B., Li, X., Zhang, Q. and Xiong, L. (2006) Overexpressing a NAM, ATAF, and CUC (NAC) transcription factor enhances drought resistance and salt tolerance in rice. Proc. Natl. Acad. Sci. U. S. A., 103, 12987–12992.

Hu, H. and Xiong, L. (2014) Genetic engineering and breeding of drought-resistant crops. Annu. Rev. Plant Biol., 65, 715–741.

Huang, D., Wang, S., Zhang, B., Shang-Guan, K., Shi, Y., Zhang, D., Liu, X., Wu, K., Xu, Z., Fu, X. and Zhou, Y. (2015) A Gibberellin-Mediated DELLA-NAC Signaling Cascade Regulates Cellulose Synthesis in Rice. Plant Cell, 27, 1681–1696.

Ishimaru, T., Ida, M., Hirose, S., Shimamura, S., Masumura, T., Nishizawa, N.K., Nakazono, M. and Kondo, M. (2015) Laser microdissection-based gene expression analysis in the aleurone layer and starchy endosperm of developing rice caryopses in the early storage phase. Rice (N Y), 8, 57.

Kawahara, Y., de la Bastide, M., Hamilton, J.P., Kanamori, H., McCombie, W.R., Ouyang, S., Schwartz, D.C., Tanaka, T., Wu, J., Zhou, S., Childs, K.L., Davidson, R.M., Lin, H., Quesada-Ocampo, L., Vaillancourt, B., Sakai, H., Lee, S.S., Kim, J., Numa, H., Itoh, T., Buell, C.R. and Matsumoto, T. (2013) Improvement of the Oryza sativa Nipponbare reference genome using next generation sequence and optical map data. Rice (N Y), 6, 4.

Kim, D., Langmead, B. and Salzberg, S.L. (2015) HISAT: a fast spliced aligner with low memory requirements. Nat. Methods, 12, 357–360.

Koini, M.A., Alvey, L., Allen, T., Tilley, C.A., Harberd, N.P., Whitelam, G.C. and Franklin, K.A. (2009) High temperature-mediated adaptations in plant architecture require the bHLH transcription factor PIF4. Curr. Biol., 19, 408–413.

Kouchi, H. and Hata, S. (1993) Isolation and characterization of novel nodulin cDNAs representing genes expressed at early stages of soybean nodule development. Mol. Gen. Genet., 238, 106–119.

Li, B., Gao, K., Ren, H. and Tang, W. (2018) Molecular mechanisms governing plant responses to high temperatures. J. Integr. Plant Biol., 60, 757–779.

Li, H. and Durbin, R. (2009) Fast and accurate short read alignment with Burrows-Wheeler transform. Bioinformatics, 25, 1754–1760.

Lim, S.D., Cho, H.Y., Park, Y.C., Ham, D.J., Lee, J.K. and Jang, C.S. (2013) The rice RING finger E3 ligase, OsHCI1, drives nuclear export of multiple substrate proteins and its heterogeneous overexpression enhances acquired thermotolerance. J. Exp. Bot., 64, 2899–2914.

Lin, M.Y., Chai, K.H., Ko, S.S., Kuang, L.Y., Lur, H.S. and Charng, Y.Y. (2014) A positive feedback loop between HEAT SHOCK PROTEIN101 and HEAT STRESS-ASSOCIATED 32-KD PROTEIN modulates long-term acquired thermotolerance illustrating diverse heat stress responses in rice varieties. Plant Physiol., 164, 2045–2053.

Lin, Y.J. and Zhang, Q. (2005) Optimising the tissue culture conditions for high efficiency transformation of indica rice. Plant Cell Rep, 23, 540–547.

Lindemose, S., Jensen, M.K., Van de Velde, J., O’Shea, C., Heyndrickx, K.S., Workman, C.T., Vandepoele, K., Skriver, K. and De Masi, F. (2014) A DNA-binding-site landscape and regulatory network analysis for NAC transcription factors in Arabidopsis thaliana. Nucleic Acids Res., 42, 7681–7693.

Liu, H., Ding, Y., Zhou, Y., Jin, W., Xie, K. and Chen, L.L. (2017) CRISPR-P 2.0: An Improved CRISPR-Cas9 Tool for Genome Editing in Plants. Mol Plant, 10, 530–532.

Liu, H.T., Li, B., Shang, Z.L., Li, X.Z., Mu, R.L., Sun, D.Y. and Zhou, R.G. (2003) Calmodulin is involved in heat shock signal transduction in wheat. Plant Physiol., 132, 1186–1195.

Liu, W., Xie, X., Ma, X., Li, J., Chen, J. and Liu, Y.G. (2015) DSDecode: A Web-Based Tool for Decoding of Sequencing Chromatograms for Genotyping of Targeted Mutations. Mol Plant, 8, 1431–1433.

Livak, K.J. and Schmittgen, T.D. (2001) Analysis of relative gene expression data using real-time quantitative PCR and the 2(-Delta Delta C(T)) Method. Methods, 25, 402–408.

Love, M.I., Huber, W. and Anders, S. (2014) Moderated estimation of fold change and dispersion for RNA-seq data with DESeq2. Genome Biol., 15.

Lu, K., Li, T., He, J., Chang, W., Zhang, R., Liu, M., Yu, M., Fan, Y., Ma, J., Sun, W., Qu, C., Liu, L., Li, N., Liang, Y., Wang, R., Qian, W., Tang, Z., Xu, X., Lei, B., Zhang, K. and Li, J. (2018) qPrimerDB: a thermodynamics-based gene-specific qPCR primer database for 147 organisms. Nucleic Acids Res., 46, D1229–D1236.

Ma, X., Zhang, Q., Zhu, Q., Liu, W., Chen, Y., Qiu, R., Wang, B., Yang, Z., Li, H., Lin, Y., Xie, Y., Shen, R., Chen, S., Wang, Z., Chen, Y., Guo, J., Chen, L., Zhao, X., Dong, Z. and Liu, Y.G. (2015) A Robust CRISPR/Cas9 System for Convenient, High-Efficiency Multiplex Genome Editing in Monocot and Dicot Plants. Mol Plant, 8, 1274–1284.

Machanick, P. and Bailey, T.L. (2011) MEME-ChIP: motif analysis of large DNA datasets. Bioinformatics, 27, 1696–1697.

Mao, C., Ding, W., Wu, Y., Yu, J., He, X., Shou, H. and Wu, P. (2007) Overexpression of a NAC-domain protein promotes shoot branching in rice. New Phytol., 176, 288–298.

Mathew, I.E., Das, S., Mahto, A. and Agarwal, P. (2016) Three Rice NAC Transcription Factors Heteromerize and Are Associated with Seed Size. Front Plant Sci, 7, 1638.

Matsuda, T., Kawahara, H. and Chonan, N. (1979) Histo-Cytological Researches on Translocation and Ripening in Rice Ovary: I. Histological changes and transfer pathways in the developing ovary. Japanese journal of crop science, 48, 155–162.

Nuruzzaman, M., Manimekalai, R., Sharoni, A.M., Satoh, K., Kondoh, H., Ooka, H. and Kikuchi, S. (2010) Genome-wide analysis of NAC transcription factor family in rice. Gene, 465, 30–44.

Olsen, A.N., Ernst, H.A., Leggio, L.L. and Skriver, K. (2005) DNA-binding specificity and molecular functions of NAC transcription factors. Plant Sci., 169, 785–797.

Ooka, H., Satoh, K., Doi, K., Nagata, T., Otomo, Y., Murakami, K., Matsubara, K., Osato, N., Kawai, J., Carninci, P., Hayashizaki, Y., Suzuki, K., Kojima, K., Takahara, Y., Yamamoto, K. and Kikuchi, S. (2003) Comprehensive analysis of NAC family genes in Oryza sativa and Arabidopsis thaliana. DNA Res., 10, 239–247.

Oono, Y., Wakasa, Y., Hirose, S., Yang, L., Sakuta, C. and Takaiwa, F. (2010) Analysis of ER stress in developing rice endosperm accumulating beta-amyloid peptide. Plant Biotechnol. J., 8, 691–718.

Oparka, K.J. and Gates, P. (1981) Transport of assimilates in the developing caryopsis of rice (Oryza sativa L.): The pathways of water and assimilated carbon. Planta, 152, 388–396.

Patrick, J.W. (1997) PHLOEM UNLOADING: Sieve Element Unloading and Post-Sieve Element Transport. Annu. Rev. Plant Physiol. Plant Mol. Biol., 48, 191222.

Qi, W., Sun, F., Wang, Q., Chen, M., Huang, Y., Feng, Y.Q., Luo, X. and Yang, J. (2011) Rice ethylene-response AP2/ERF factor OsEATB restricts internode elongation by down-regulating a gibberellin biosynthetic gene. Plant Physiol., 157, 216–228.

Robinson, J.T., Thorvaldsdottir, H., Winckler, W., Guttman, M., Lander, E.S., Getz, G. and Mesirov, J.P. (2011) Integrative genomics viewer. Nat. Biotechnol., 29, 24–26.

Schramm, F., Larkindale, J., Kiehlmann, E., Ganguli, A., Englich, G., Vierling, E. and von Koskull-Doring, P. (2008) A cascade of transcription factor DREB2A and heat stress transcription factor HsfA3 regulates the heat stress response of Arabidopsis. Plant J., 53, 264–274.

Shen, J., Liu, J., Xie, K., Xing, F., Xiong, F., Xiao, J., Li, X. and Xiong, L. (2017a) Translational repression by a miniature inverted-repeat transposable element in the 3’ untranslated region. Nat Commun, 8, 14651.

Shen, J., Lv, B., Luo, L., He, J., Mao, C., Xi, D. and Ming, F. (2017b) The NAC-type transcription factor OsNAC2 regulates ABA-dependent genes and abiotic stress tolerance in rice. Sci. Rep., 7, 40641.

Singh, A., Mittal, D., Lavania, D., Agarwal, M., Mishra, R.C. and Grover, A. (2012) OsHsfA2c and OsHsfB4b are involved in the transcriptional regulation of cytoplasmic OsClpB (Hsp100) gene in rice (Oryza sativa L.). Cell Stress Chaperones, 17, 243–254.

Sosso, D., Luo, D., Li, Q.B., Sasse, J., Yang, J., Gendrot, G., Suzuki, M., Koch, K.E., McCarty, D.R., Chourey, P.S., Rogowsky, P.M., Ross-Ibarra, J., Yang, B. and Frommer, W.B. (2015) Seed filling in domesticated maize and rice depends on SWEET-mediated hexose transport. Nat. Genet., 47, 1489–1493.

Waadt, R., Schmidt, L.K., Lohse, M., Hashimoto, K., Bock, R. and Kudla, J. (2008) Multicolor bimolecular fluorescence complementation reveals simultaneous formation of alternative CBL/CIPK complexes in planta. Plant J., 56, 505–516.

Walter, M., Chaban, C., Schutze, K., Batistic, O., Weckermann, K., Nake, C., Blazevic, D., Grefen, C., Schumacher, K., Oecking, C., Harter, K. and Kudla, J. (2004) Visualization of protein interactions in living plant cells using bimolecular fluorescence complementation. Plant J., 40, 428–438.

Wang, L., Xie, W., Chen, Y., Tang, W., Yang, J., Ye, R., Liu, L., Lin, Y., Xu, C., Xiao, J. and Zhang, Q. (2010) A dynamic gene expression atlas covering the entire life cycle of rice. Plant J., 61, 752–766.

Wang, L., Zhou, Z., Song, X., Li, J., Deng, X. and Mei, F. (2008a) Evidence of ceased programmed cell death in metaphloem sieve elements in the developing caryopsis of Triticum aestivum L. Protoplasma, 234, 87–96.

Wang, Y., Xiao, Y., Zhang, Y., Chai, C., Wei, G., Wei, X., Xu, H., Wang, M., Ouwerkerk, P.B. and Zhu, Z. (2008b) Molecular cloning, functional characterization and expression analysis of a novel monosaccharide transporter gene OsMST6 from rice (Oryza sativa L.). Planta, 228, 525–535.

Wu, X., Liu, J., Li, D. and Liu, C.M. (2016) Rice caryopsis development II: Dynamic changes in the endosperm. J. Integr. Plant Biol., 58, 786–798.

Xiong, Y., Ren, Y., Li, W., Wu, F., Yang, W., Huang, X. and Yao, J. (2019) NF-YC12 is a key multi-functional regulator of accumulation of seed storage substances in rice. J. Exp. Bot., 70, 3765–3780.

Xu, G.Y., Rocha, P.S., Wang, M.L., Xu, M.L., Cui, Y.C., Li, L.Y., Zhu, Y.X. and Xia, X. (2011) A novel rice calmodulin-like gene, OsMSR2, enhances drought and salt tolerance and increases ABA sensitivity in Arabidopsis. Planta, 234, 47–59.

Xue, W., Xing, Y., Weng, X., Zhao, Y., Tang, W., Wang, L., Zhou, H., Yu, S., Xu, C., Li, X. and Zhang, Q. (2008) Natural variation in Ghd7 is an important regulator of heading date and yield potential in rice. Nat. Genet., 40, 761–767.

Yamaguchi, M., Ohtani, M., Mitsuda, N., Kubo, M., Ohme-Takagi, M., Fukuda, H. and Demura, T. (2010) VND-INTERACTING2, a NAC domain transcription factor, negatively regulates xylem vessel formation in Arabidopsis. Plant Cell, 22, 1249–1263.

Yang, J., Luo, D., Yang, B., Frommer, W.B. and Eom, J.S. (2018) SWEET11 and 15 as key players in seed filling in rice. New Phytol.

Yang, S.D., Seo, P.J., Yoon, H.K. and Park, C.M. (2011) The Arabidopsis NAC transcription factor VNI2 integrates abscisic acid signals into leaf senescence via the COR/RD genes. Plant Cell, 23, 2155–2168.

Young, M.D., Wakefield, M.J., Smyth, G.K. and Oshlack, A. (2010) Gene ontology analysis for RNA-seq: accounting for selection bias. Genome Biol., 11, R14.

Zhang, Y., Liu, T., Meyer, C.A., Eeckhoute, J., Johnson, D.S., Bernstein, B.E., Nusbaum, C., Myers, R.M., Brown, M., Li, W. and Liu, X.S. (2008) Model-based analysis of ChIP-Seq (MACS). Genome Biol., 9, R137.

Zhang, Z., Dong, J., Ji, C., Wu, Y. and Messing, J. (2019) NAC-type transcription factors regulate accumulation of starch and protein in maize seeds. Proc. Natl. Acad. Sci. U. S. A., 116, 11223–11228.

