## Supplemental Figures for "A Heat Stress Responsive NAC Transcription Factor Heterodimer Plays Key Roles in Rice Grain Filling"

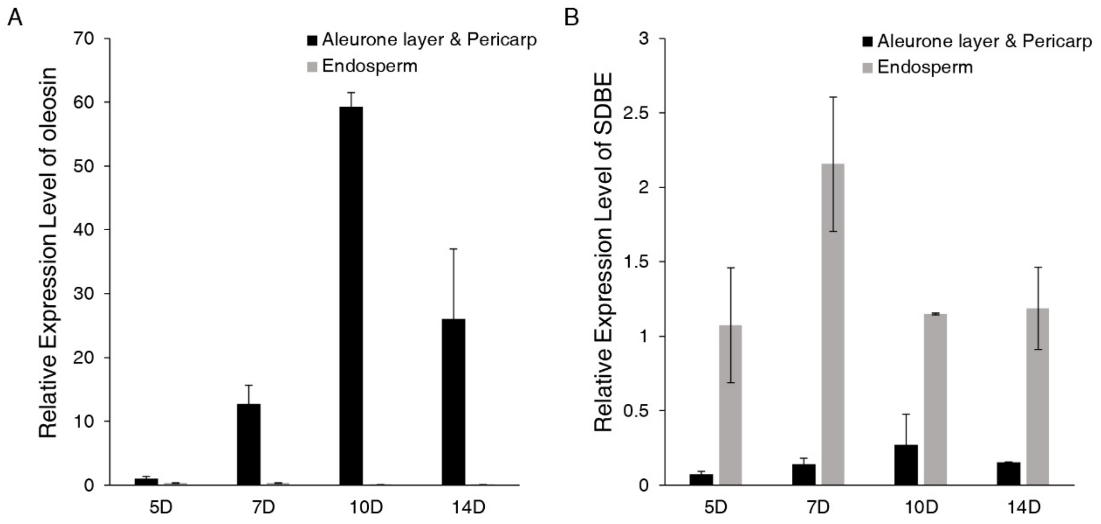


**Supplementary Fig. S1. Expression levels of the markers for mechanical isolation of different tissues in rice seeds.** (a) Aleurone layer marker *oleosin*. (b) Endosperm marker *SDBE*. 5-14D, total RNA was extracted from seeds collected during 5–14 DAP. The values in each column are the mean of three technical replicates and the error bars indicate the ±SD. Ubiquitin was used as the reference gene.

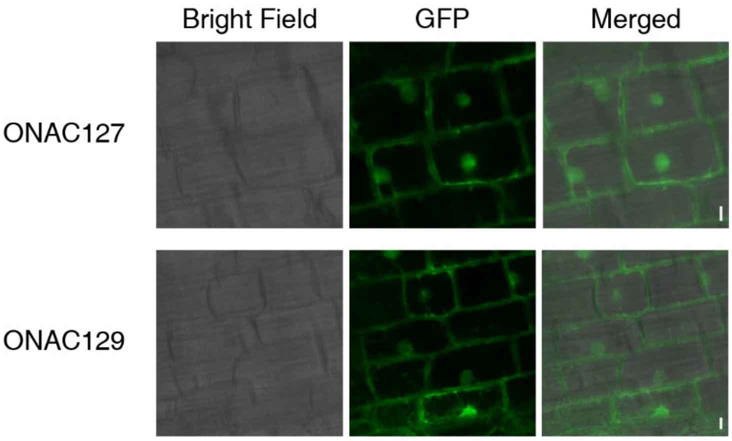


**Supplementary Fig. S2. Subcellular localization of ONAC127 and ONAC129 in the roots of two-week-old rice seedlings of pUbi::ONAC127-GFP and pUbi::ONAC129-GFP transgenic plants.** Scale bars = 5 μm.

| | ONAC127 | | ONAC129 |
|---|---|---|---|
| *onac127-1* | CCTTGG– –TTCCACTACGGGAAT | *onac129-1* | CCCTCTAGAATTCATTTGTGATGT |
| ZH11 | CCTTGGTATTCCACTACGGGAAT | ZH11 | CCCTCT–GAATTCATTTGTGATGT |
| *onac127-2* | CCTTGGTTATTCCACTACGGGAAT | *onac129-2* | ACGTTCTTCCTCATA– –CCATGG |
| ZH11 | CCTTGG–TATTCCACTACGGGAAT | ZH11 | ACGTTCTTCCTCATAACCCATGG |
| *onac127-3* | CCCTGTGGACTTCATCACCAACGT | *onac129-3* | ACGTTCTTCCTCATAA–CCATGG |
| ZH11 | CCCTGT–GACTTCATCACCAACGT | ZH11 | ACGTTCTTCCTCATAACCCATGG |

| | | | |
|---|---|---|---|
| *onac127;129-1* | CCGTGC– – – – – – –GATGGGTTCT | *onac127;129-1* | GGGCATGTCACACCC– –CCGAGG |
| ZH11 | CCGTGCTGCTGGTGATGGGTTCT | ZH11 | GGGCATGTCACACCCCACCGAGG |
| *onac127;129-2* | CCGTGC–GCTGGTGATGGGTTCT | *onac127;129-2* | GGGCATGTCACACCCCAACCGAGG |
| ZH11 | CCGTGCTGCTGGTGATGGGTTCT | ZH11 | GGGCATGTCACACCCC–ACCGAGG |
| *onac127;129-3* | CCCTG– – – –TTCATCACCAACGTC | *onac127;129-3* | CCCCGC– –TGGGCATGTCACACC |
| ZH11 | CCCTGTGACTTCATCACCAACGTC | ZH11 | CCCCGCTGTGGGCATGTCACACC |

**Supplementary Fig. S3. Mutation sites in *onac127*, *onac129* and *onac127;129* lines as compared with wild-type (ZH11) sequences.** The protospacer-adjacent motif sequences are shown in orange and the inserted or deleted nucleotides are indicated in red.

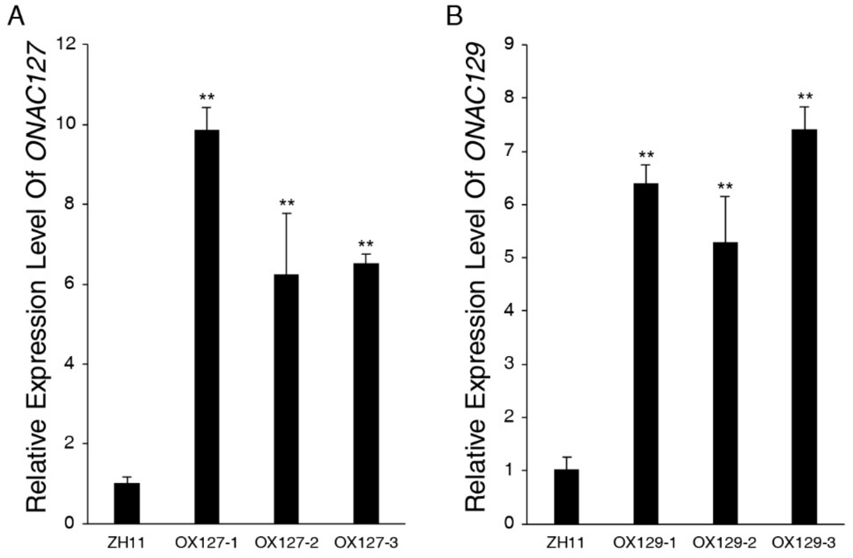


**Supplementary Fig. S4. Relative expression levels of *ONAC127* and *ONAC129* in the overexpression lines.** The values in each column are the mean (±SD) of three biological replicates. Ubiquitin was used as the reference gene. Significant differences between the ZH11 and the overexpression lines were determined using Student's t-test (** $P$<0.01).

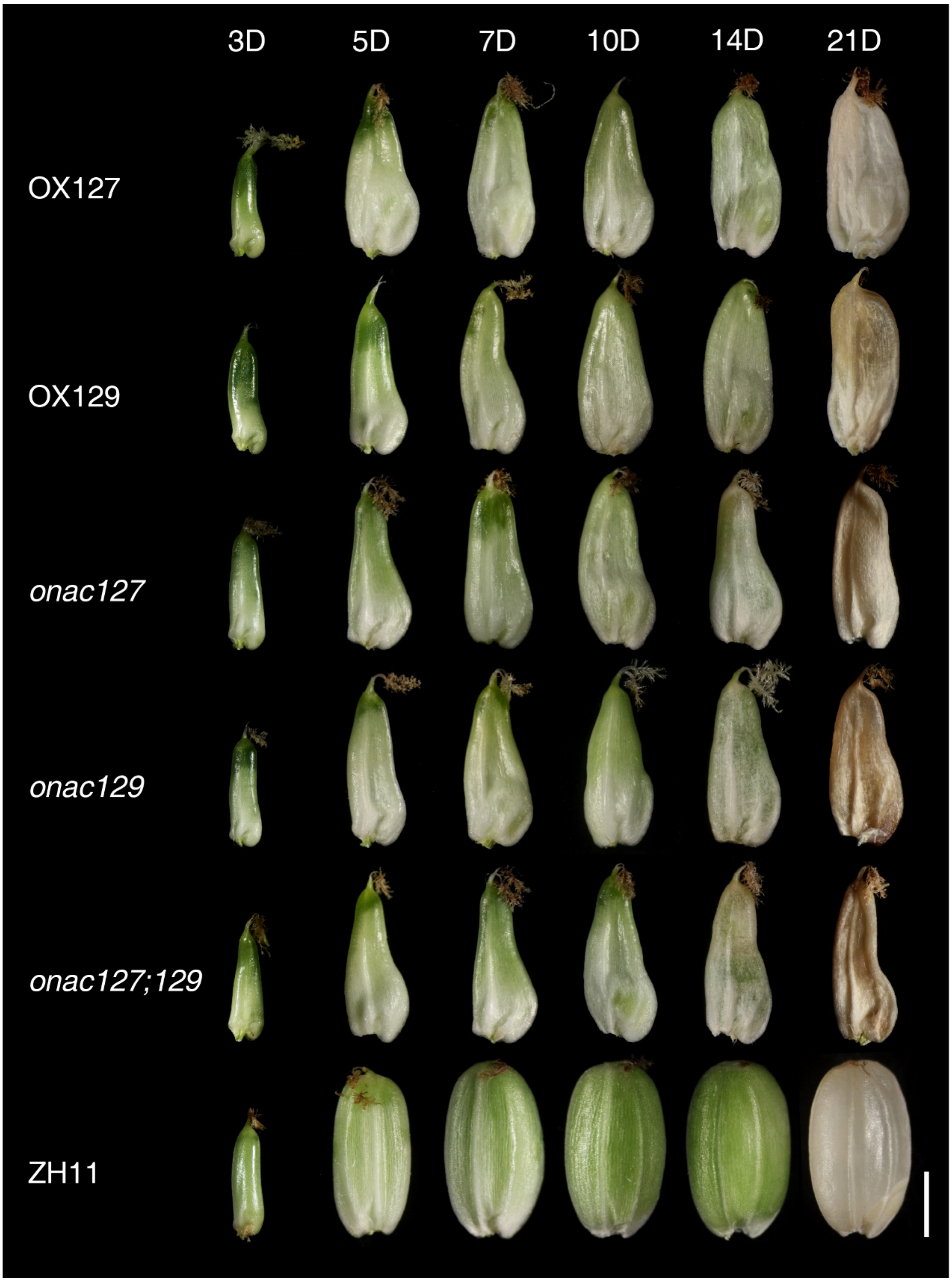


**Supplementary Fig. S5**. **Different stages of seed development.** Scale bar = 2 mm.

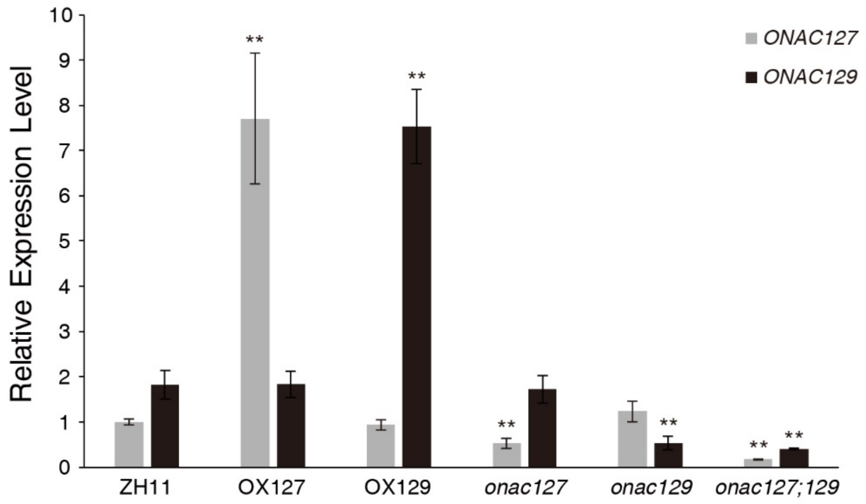


**Supplementary Fig. S6. Relative expression levels of *ONAC127* and *ONAC129* in 7-DAP seeds of transgenic lines compared with that of ZH11.** Ubiquitin was used as the reference gene. The values in each column are the mean (±SD) of three replicates. Significant differences between the ZH11 and the transgenic lines were determined using Student's t-test (** $P$<0.01).

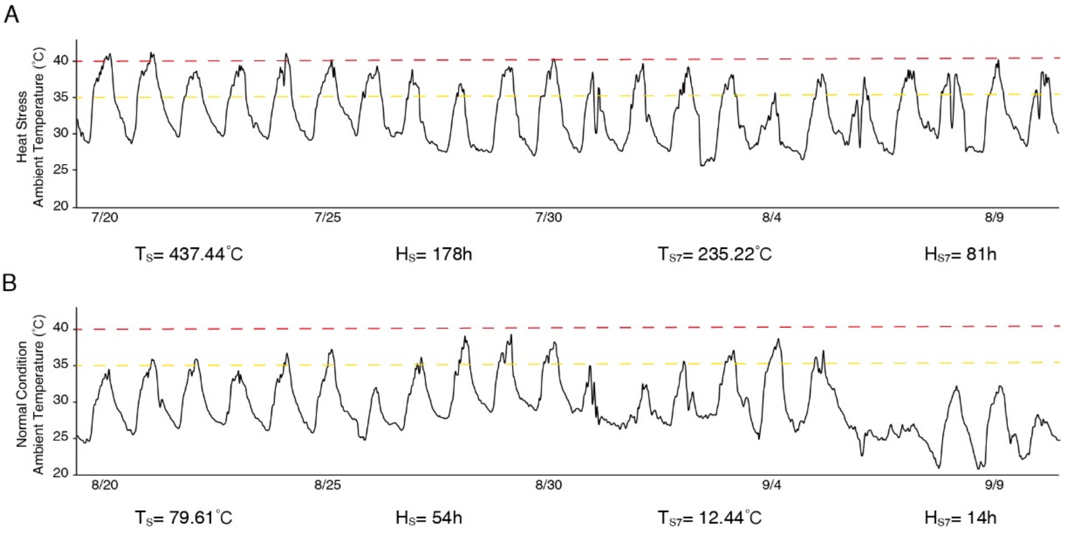


**Supplementary Fig. S7. Ambient temperature of growing areas during rice grain filling stage. 35℃ is marked as yellow dotted lines and 40℃ is marked as red dotted lines.** $T_S$, heat damage accumulated temperature during the whole filling stage; $H_S$, heat damage hours during the whole filling stage; $T_{S7}$, heat damage accumulated temperature during 0–7 DAP; $H_{S7}$, heat damage hours during 0–7 DAP.

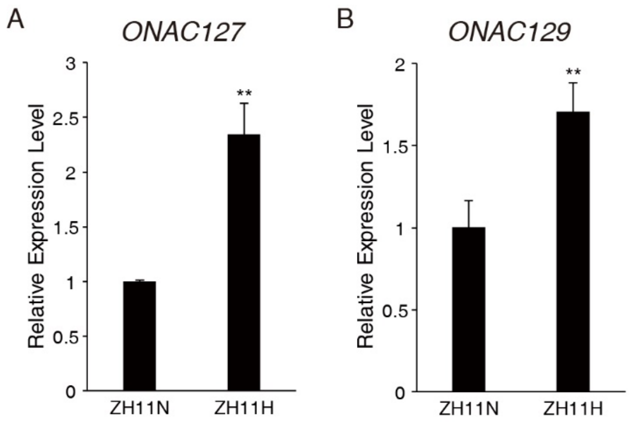


**Supplementary Fig. S8. Relative expression levels of *ONAC127* and *ONAC129* in 7-DAP seeds of ZH11 under heat stress and normal conditions.** ZH11H represents the sample collected under heat stress, and ZH11N represents the sample collected under normal conditions. Ubiquitin was used as the reference gene. The values in each column are the mean (±SD) of three replicates. Significant differences between the samples were determined using Student's t-test (** $P$<0.01).

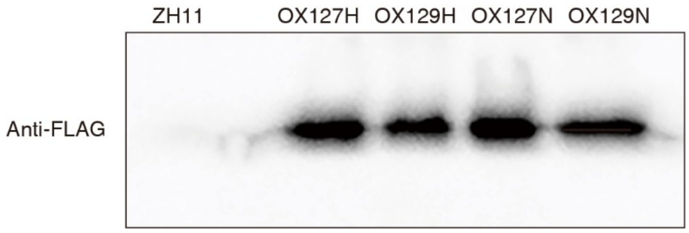


**Supplementary Fig. S9. Detection of FLAG fusion proteins in ZH11 and overexpression lines.** Total proteins extracted from developing caryopses at 7-DAP were used for western blot analysis with an anti-FLAG antibody.

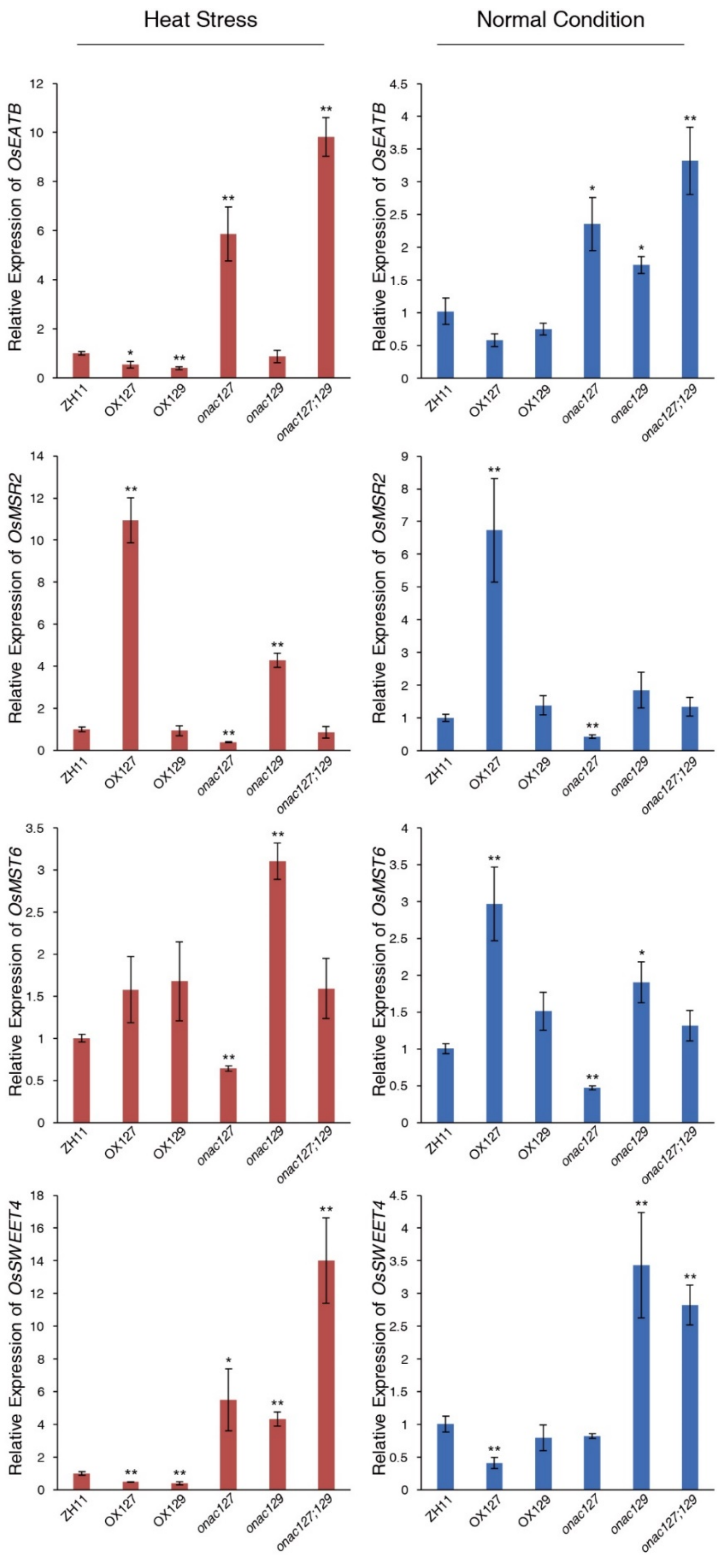


**Supplementary Fig. S10. Expression levels of the target genes of ONAC127 and ONAC129 in 7-DAP seeds of transgenic lines compared with that of ZH11.** Ubiquitin was used as the reference gene. The values in each column are the mean (±SD) of three replicates. Significant differences between the ZH11 and the overexpression lines were determined using Student's t-test (* $P<0.05$; ** $P<0.01$).

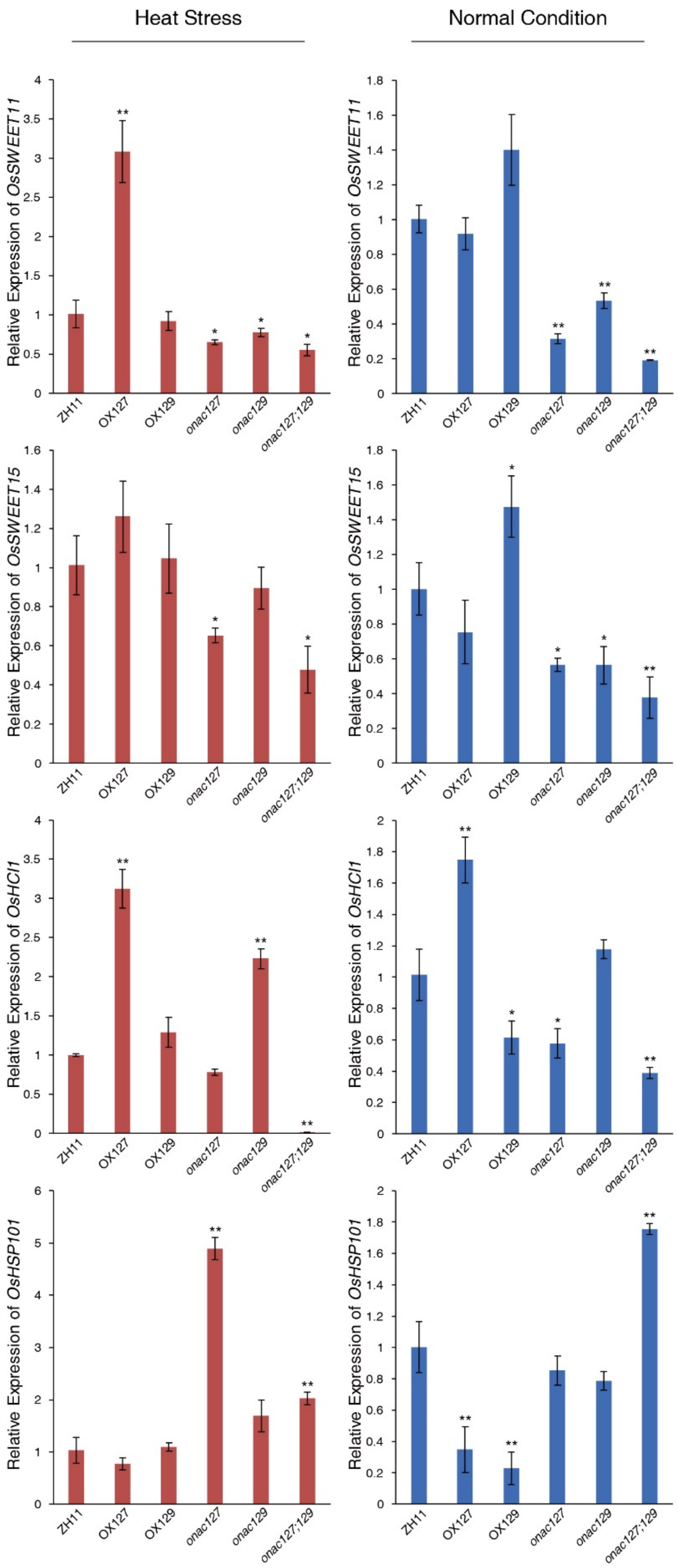


**Supplementary Fig. S11. Expression levels of the indirect target genes of ONAC127 and ONAC129 in 7-DAP seeds of transgenic lines compared with that of ZH11.** Ubiquitin was used as the reference gene. The values in each column are the mean (±SD) of three replicates. Significant differences between ZH11 and the overexpression lines are determined using Student's t-test (* $P$<0.05; ** $P$<0.01).
